# Long-range alpha-synchronisation as control signal for BCI: A feasibility study

**DOI:** 10.1101/2022.05.25.493035

**Authors:** Martín Esparza-Iaizzo, Irene Vigué-Guix, Manuela Ruzzoli, Mireia Torralba, Salvador Soto-Faraco

**Affiliations:** Center for Brain and Cognition, Universitat Pompeu Fabra, Barcelona, Spain; Basque Center on Cognition Brain and Language (BCBL), Donostia/San Sebastián, Spain; Ikerbasque, Basque Foundation for Science, Bilbao, Spain; Institució Catalana de Recerca i Estudis Avançats (ICREA), Barcelona, Spain

**Author notes:** Corresponding author: Salvador Soto-Faraco, Universitat Pompeu Fabra, Room 24.327, Ed. Mercè Rodoreda, Carrer de Ramón Trias Fargas, 25-27, 08005 Barcelona, Spain.

**Keywords:** Brain-Computer Interface, Alpha, EEG, Oscillations, Visuospatial Attention, Phase Coupling, Support Vector Machine

## Abstract

Shifts in spatial attention are associated with variations in alpha-band (α, 8–14 Hz) activity, specifically in inter-hemispheric imbalance. The underlying mechanism is attributed to local α-synchronisation, which regulates local inhibition of neural excitability, and fronto-parietal synchronisation reflecting long-range communication. The direction-specific nature of this neural correlate brings forward its potential as a control signal in brain-computer interfaces (BCI). In the present study, we explored whether long-range α-synchronisation presents lateralised patterns dependent on voluntary attention orienting and whether these neural patterns can be picked up at a single-trial level to provide a control signal for active BCI. We collected electroencephalography (EEG) data from a cohort of healthy adults (n = 10) while performing a covert visuospatial attention (CVSA) task. The data shows a lateralised pattern of α-band phase coupling between frontal and parieto-occipital regions after target presentation, replicating previous findings. This pattern, however, was not evident during the cue-to-target orienting interval, the ideal time window for BCI. Furthermore, decoding the direction of attention trial-by-trial from cue-locked synchronisation with support vector machines (SVM) was at chance-level. The present findings suggest EEG may not be capable of detecting long-range α-synchronisation in attentional orienting on a single-trial basis and, thus, highlight the limitations of this metric as a reliable signal for BCI control.

**SIGNIFICANCE STATEMENT:** Cognitive neuroscience advances should ideally have a real-world impact, with an obvious avenue for transference being BCI applications. The hope is to faithfully translate user-generated brain endogenous states into control signals to actuate devices. A paramount challenge for transfer is to move from group-level, multi-trial average approaches to single-trial level. Here, we evaluated the feasibility of single-trial estimation of phase synchrony across distant brain regions. Although many studies link attention to long-range synchrony modulation, this metric has never been used to control BCI. We present a first attempt of a synchrony-based BCI that, albeit unsuccessful, should help break new ground to map endogenous attention shifts to real-time control of brain-computer actuated systems.

## INTRODUCTION

A few decades ago, imagining an interface between the human brain and a computer was closer to science fiction than to scientific achievement. Nowadays, brain-computer interfaces (BCIs) can read out brain activity, extract features from the signal in real-time, and convert them into outputs for monitoring, controlling devices, or even modifying cognitive states (Blankertz et al. 2016). One significant challenge of BCIs is finding reliable control signals from brain activity with a sufficiently high signal-to-noise ratio (SNR) at a trial-by-trial level to allow successful classification. Ideally, the appearance of the target brain activity should depend on endogenous mental states that a user can control at will. The use of non-invasive, cost-effective, and light-weight neuroimaging devices can, in turn, facilitate transfer to real applications. For now, EEG is the most viable candidate to achieve real-life BCI.

For example, some EEG-based BCIs have used motor imagery as a control signal (e.g., imagined right/left-limb movement; Padfield et al. 2019), whereas others have used neural correlates of covert visuospatial attention (CVSA) (van Gerven and Jensen 2009; Treder et al. 2011; Tonin et al. 2013). Here, we will concentrate on the latter. In human behaviour, CVSA is used to direct processing resources to relevant locations in the environment whilst disengaging from irrelevant locations (Pashler 1999; Foster and Awh 2019). CVSA can be manipulated through a Posner cueing protocol (Posner 1980), which shows a robust effect on behavioural performance: higher accuracy and faster reaction times for targets appearing at the cued (attended) location compared to targets appearing in un-cued, putatively unattended locations (Posner 1980).

Shifts in CVSA are associated with changes in oscillatory activity in the alpha-band (α, 8–14 Hz) at parieto-occipital regions (Klimesch 1999; Foster et al. 2017). Typically, α-power shows an inter-hemispheric imbalance when attention is covertly oriented to either the left or right visual field, revealing its potential as a control signal for BCI implementations (Rihs et al. 2007; Thut et al. 2006, see Astrand et al. 2014b for a review). Inter-hemispheric α-power imbalance corresponds to a late process in CSVA shifts (van Diepen et al. 2019). First, cueing information is integrated through sensory pathways in a bottom-up fashion, reaching higher visual areas in the parietal cortex (e.g., intraparietal sulcus) and eventually frontal regions (e.g., frontal eye fields) (Petersen and Posner 2012). From there on, top-down modulation shifts attention to the corresponding hemifield, where it is maintained during target anticipation (Simpson et al., 2012). The mechanism involved in this top-down modulation is thought to involve long-range α-synchronisation between the frontal and posterior cortex, which eventually leads to classical inter-hemispheric imbalances in α-power observed in the visual cortex (Sauseng et al. 2005; Doesburg et al. 2009, Lobier et al. 2018). Long-range synchronisation is a potential mechanism to increase the fidelity and effectiveness of communication throughout the brain (Clayton et al. 2018) among occipital, parietal, and frontal regions (Sadaghiani and Kleinschmidt 2016). Synchronising excitability cycles between distant neural populations increases the likelihood of spikes from one region discharging post-synaptic potentials during a specific (excitable) phase of the other (Fries, 2015). Despite the evidence supporting this model (Buschman and Miller., 2007; Cardin et al., 2009), there is still debate on its temporal dynamics, lateralisation patterns and individual-level variability.

Despite the evidence of links between long-range α-synchronisation and behavioural performance at group-level analyses (Sauseng et al. 2005; Doesburg et al. 2009; Doesburg et al. 2016), BCI protocols based on endogenous attention orienting have only used α-power as a control signal. In our study, we attempt to replicate a previously demonstrated effect in attention orienting involving long-range α-synchronisation to assess its feasibility in BCI paradigms. The original publication (Sauseng et al., 2005) found significant increases in contralateral over ipsilateral connectivity around the time of target appearance. We hypothesized that, if attention-driven connectivity emerged in target-centered time windows, it may also be present in the cue-to-target interval, where participants are putatively shifting attention towards the cued side. Further, this cue-to-target time window would enable the use of long-range α-synchronisation in BCIs based on purely endogenous brain signals. Therefore, we will test whether such contra- and ipsilateral patterns in α-synchronisation emerge in single-trial dynamics with sufficient signal strength to make them a reliable control signal. To do so, we used an EEG dataset from a lateralised endogenous spatial attention task to replicate group-level effects found by Sauseng et al. (2005), to explore the cue-to-target interval, and to classify the direction of attention at the single-trial level using long-range α-phase synchronisation as proof of concept for transference to BCI.

## MATERIALS AND METHODS

### Participants

We used data from a previous, unrelated study (Torralba et al 2016). The dataset consisted of 15 participants (mean age = 22, SD = 3; 7 female). All participants provided informed consent and had a normal or corrected-to-normal vision. The study was ran in accordance with the Declaration of Helsinki and the experimental protocol approved by the local ethics committee CEIC Parc de Salut Mar (Barcelona, Spain).

### Task

Before the experimental session, the participant’s EEG activity was recorded during a five-minute recording at rest with eyes closed to extract the individual α frequency (IAF) used in the analyses. In the experimental session, participants performed a modified version of the Posner cueing task (see **Figure 1A**). The trial started with the onset of a central fixation cross, placed between two placeholders located 20° of visual angle left and right off-centre, vertically shifted 20° of visual angle below the fixation cross (see **Figure 1A**). After 200 ms fixation period, a central auditory cue (100 ms duration) indicated the likely target location through either high pitch (2000 Hz) or low pitch (500 Hz) tones, the mapping was randomised across subjects. Participants should covertly attend to the indicated side, without moving their eyes, during a jittered inter-stimulus interval (ISI; 2000 ± 500 ms). The use of a jittered ISI was employed in order to avoid participants using automatic temporal attention to solve the task. Next, the target (a Gabor grating tilted 45° left or right, 50 ms duration) appeared briefly inside one of the placeholders, with 75% validity regarding the cued location. The grating contrast was adjusted individually, as described below. A noise pattern with an equal overall luminance as the target was presented at the alternative placeholder, with the exact timings as the target. Participants were asked first to indicate if they had detected the target (yes/no detection) and, subsequently, the target’s tilt (left/right discrimination). Both answers were made by keypress, in an un-speeded fashion, and with response mapping (top-bottom) orthogonal to the attention manipulation and varied from trial to trial. A trial was considered correctly answered only when participants both detected the stimulus and discriminated the hemifield in which it was presented. An inter-trial interval of 1000 ms followed the response, and a new trial began. Unless otherwise noted, the EEG analyses were done on validly cued trials that responded correctly. On average, 289.9 ± 11.3 trials from each participant were employed for the EEG analysis.

**Figure 1.**
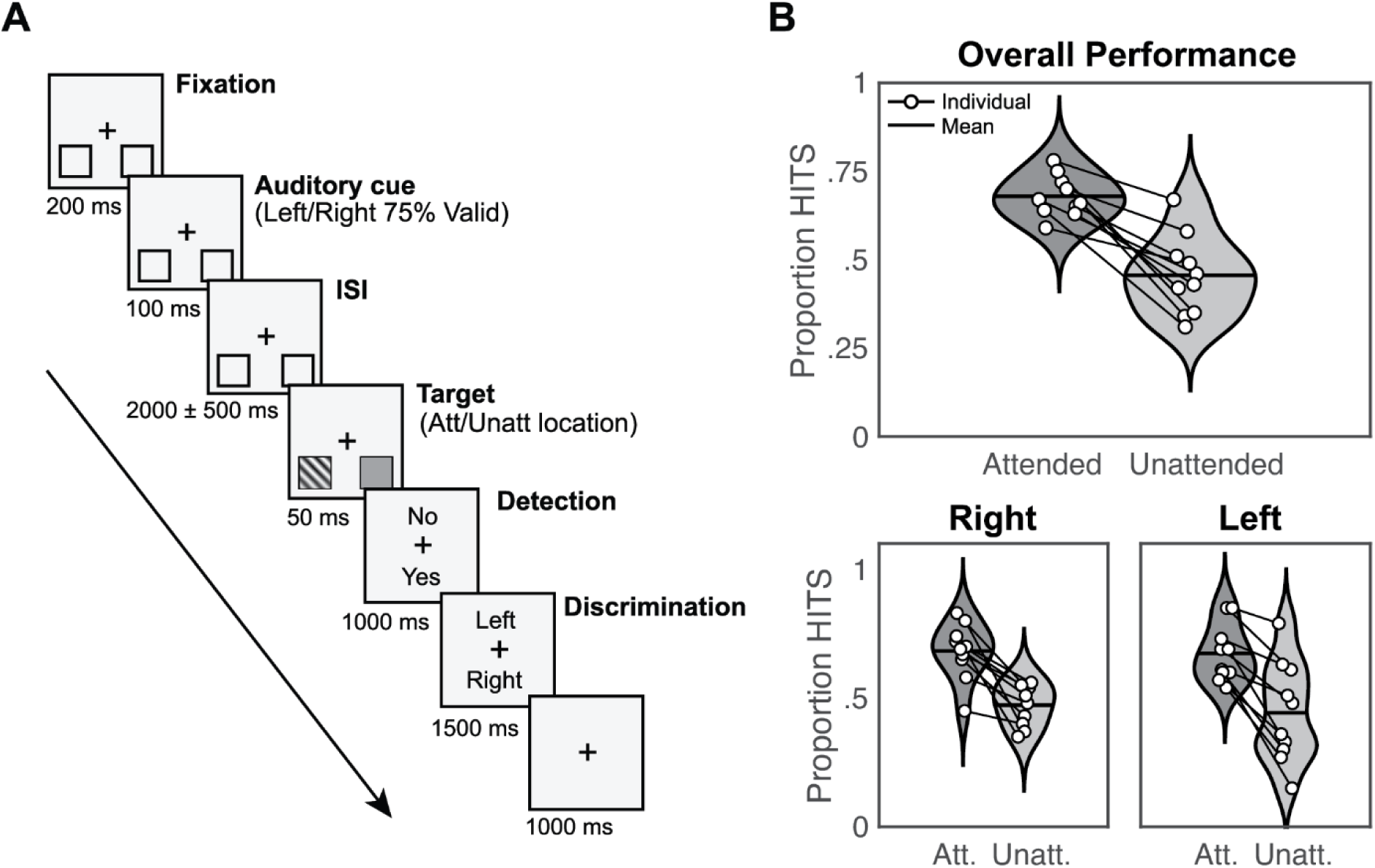
Experimental design and response rates. **(A) Schematic trial representation.** A black fixation cross in the middle of the screen and two squares (to-be-attended locations) at the bottom left, and bottom right positions were displayed continuously. At the beginning of each trial, participants were instructed to gaze at the fixation cross. After 200 ms (fixation period), an auditory cue appeared for 100 ms (cue period) indicating which hemifields participants must attend (75% validity). After a jittered interstimulus interval of 2000 ± 500 ms, a target appeared at the targeted location during 50 ms (target period). Participants had to report first if they had seen the target (detection task), and after 1000 ms, the location of the target (left/right discrimination task) during 1500 ms. An intertrial interval (ITI) of 1000 ms followed, and a new trial began (Adapted from Torralba et al. 2016). **(B) Response rates for detected and discriminated trials (HITS) related to attended and unattended trials.** Black lines over violin plots represent the mean value. Both overall performance (top) and right/left hemifields (bottom) are shown. White dots indicate individual values (adapted from Torralba et al. 2016).

The Gabor gratings used as stimuli were 0.002 cycles per degree, with a size of 3.35°, embedded in white noise. The contrast was adjusted individually using a preliminary threshold titration procedure in which thresholds for both sides (left and right) were independently adjusted to a 70% detection rate when cued (in the attended location). Stimuli were presented on a 21” CRT screen with a refresh rate of 100 Hz and a resolution of 1024 x 768 pixels. The experiment was implemented in MATLAB R2015b (MATLAB, RRID: SCR_001622) using the Psychophysics Toolbox (Psychophysics Toolbox, RRID: SCR_002881).

### EEG recording and pre-processing

EEG recordings were obtained from 64 Ag/AgCl electrodes positioned according to the 10-10 system with AFz as ground and nose tip as reference. Impedance was kept below 10 kΩ. The employed system was an active actiCHAmp EEG amplifier from Brain Products (Munich, Germany). The signal was sampled at 500 Hz and processed in MATLAB 2020 and 2015 (MATLAB, RRID: SCR_001622) using custom functions and the FieldTrip toolbox (FieldTrip, RRID: SCR_004849).

Manual artefact rejection was applied to discard trials where any EOG components had an amplitude higher than 50 µV. Defective channels were repaired using neighbours calculated by triangulation and splines for interpolating channel data. Then, the data was demeaned and notch filtered at 50 Hz to exclude line noise. Next, fifth-order high-pass and sixteenth-order low-pass IIR Butterworth filters were employed to limit the signal between 0.16 and 45 Hz (Sauseng et al. 2005). The filtering was done forward and backwards (two-pass), which resulted in zero phase lag.

### Time-frequency analysis

We performed long-range synchronisation analyses in two time windows. The first was time-locked to the target onset (target-locked) to replicate Sauseng et al. (2005) methods and validate our analysis pipeline. The second was time-locked to the cue onset (cue-locked) to estimate long-range α-phase synchronisation during covert visuospatial attention shifts.

Following Sauseng et al. (2005), for the target-locked analysis, we used two windows of 200 ms: a pre-target (-200 to 0 ms) and a post-target window (200 ms to 400 ms). The latter excludes the interval 0 to 200 ms, most affected by the phase resetting effect of target presentation. For the cue-locked analysis, we used the cue-to-target time window between 500 ms and 1500 ms post-cue and divided it into five consecutive and non-overlapping 200 ms windows. By analysing from 500 ms onwards we avoid the event related potential (ERP) caused by cue presentation and allow endogenous attention shift to build up, a process which takes a few hundreds of milliseconds (Foxe and Snyder 2011). The cue-locked analysis period ends at 1500 ms, which was the minimum possible duration of the cue-to-target interval (duration of 2000 ± 500 ms, see methods). All epoched data was mirror-reflected to avoid edge artefacts (Cohen 2014) when performing the time-frequency analysis. Afterwards, data were trimmed, and reflected edges were removed.

We computed the Fourier coefficients using 5-cycle Morlet wavelets (Grossmann and Morlet 1984) with 16 logarithmically spaced frequencies ranging from 2.6 Hz to 42 Hz. For the analysis aimed at replicating Sauseng’s results, we only used wavelets within the upper α-band (9.54 – 14.31 Hz) (Sauseng et al. 2005), whereas, for the exploratory analysis, we used the whole frequency range (i.e., 2.6 – 42 Hz) to explore further long-range α-phase synchronisation in other frequency bands beyond the IAF.

### Connectivity measures

Three clusters of electrodes of interest (EOI) were defined for the connectivity analyses, mimicking Sauseng et al., 2005: A fronto-medial (FM) EOI cluster (Fz, FC1, FC2) and two symmetric posterior clusters located either atthe parietal left (PL) region (P3, PO3, PO1) or the parietal right (PR) region (P4, PO4, PO2). To infer connectivity between each parietal EOI cluster and the FM location, we used Phase Locking Value (PLV) (Lachaux et al. 1999). This metric reports the consistency of phase differences between two locations across multiple trials and is not affected by power differences. Mathematically, the PLV is expressed as the absolute value of the average complex unit-length phase differences:

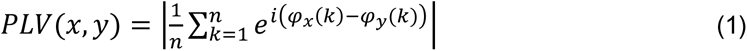

where *n* corresponds to the total number of trials indexed by *k* and *φ_x_*, *φ_y_* correspond to the phases at electrodes *x* and *y,* respectively. PLV was calculated according to equation (1) using the phases for every combination of individual electrode pairs of the FM-PR and FM-PL networks. Then, these values were averaged, resulting in a time series of PLV FM-PR and FM-PL networks for each of the frequencies of interest and condition (*attended left* and *attended right)* trials. Subsequently, the PLV time series were collapsed as either ipsilateral (FM-PL network and attend left; FM-PR and attend right) or contralateral (FM-PR network and attend left; FM-PL and attend right). Therefore, for each participant and frequency of interest, two time series of PLV were obtained (contra- and ipsilateral PLV).

### Classification

The trial classification was performed using support vector machines (SVM). We selected FM-PR and FM-PL connectivity as input to the SVM. *Attended right* and *attended left* labels for each trial were provided as ground truth for the algorithm. The main goal of the classifier was to infer, on each trial, whether a participant was attending on the left or right hemifield, based on the long-range α-phase synchronisation in the left and right fronto-parietal networks. Note that PLV is computed across trials, and SVM aims to classify on a single-trial basis, so PLV was also calculated across time points (Cohen 2015). As a validation step, we repeated the target-locked analysis employing this metric (i.e., cross-time PLV) before proceeding with the cue-locked classification attempt.

We divided the cue-locked interval ranging from 500 to 1500 ms in bins of 200 ms, yielding five values for FM-PR connectivity and five for FM-PL connectivity. The resulting ten values were used as input to the SVM to perform the optimisation and classification of the trials. Note that for the classification, we used the data from the participant that achieved a significant difference in PLV values between parietal left and right EOI clusters in all cue-to-target windows (P10). Trials were split into a training (80%) and testing (20%) set of trials to avoid overfitting. Then, the training set was subdivided into sub-training (80%) and validation sets (20%).

Our initial approach was to use a linear kernel for the classification. However, after evaluating the option through cross-validation of the validation set and obtaining a negative result (i.e., classification was not better than chance level), we decided to use a Gaussian kernel (i.e., Radial Basis Function). In order to select the most suitable and efficient values for classifying *attended left* and *attended right* trials from the validation set, we optimised the parametric space of the SVM. This comprised margin and gamma (γ) parameters, which were explored in logarithmic steps from 10^-6^ to 10^3^ for both constants and every fold.

### Inter-hemispheric power imbalance analysis

Besides calculating the long-range α-phase coupling, we also computed the inter-hemispheric α-power imbalance at parietal regions, both at the individual and at group-level, as a reality check. For this reality check, we used Thut et al., (2006) for guidance to choose the electrodes of interest. First, we performed an independent component analysis (ICA), during which 3±1 components were discarded on average per participant, based on a visual inspection, the components’ topography, and time course. The rejected components comprised both ocular and motor artifacts. Please note that ICA was only performed for the power analysis, not for the connectivity pipeline, in order to replicate the exact pre-processing as seen in Sauseng et al., (2005) and, importantly, because phase of electrophysiological recordings is affected when ICA are rejected (Thatcher et al., 2020).

The frequency of interest used in lateralisation analyses was adjusted for each participant depending on the individual α frequency (IAF) extracted from the five-minute recording (eyes closed) previous to the experiment (see above). The IAF was determined based on the presence of a single peak (i.e., a local maximum) within the considered frequency band of interest (5-15 Hz) on the power spectrum density (PSD). A spectrogram was extracted for each parieto-occipital electrode (P7, P5, P3, P1, Pz, P2, P4, P6, P8, PO3, PO4, POz, PO9, PO10, O1, Oz, O2) using the Welch method (segments of 1000 ms with a 10% overlap, a Hanning taper to avoid spectral leakage and 0.25 Hz frequency resolution). The power spectrum was averaged across electrodes for each participant and normalised by the mean power from 1 to 40 Hz (Vigué-Guix et al., 2022).

To extract the α-power during the task, we selected the epoch from -1.5 to 3 s in cue-locked trials by convolving the EEG signal with a set of complex Morlet wavelets (Grossmann and Morlet 1984) of 5 cycles (n_c_). The frequencies of the wavelets ranged from IAF ± 1 Hz, in 1 Hz steps. For instance, an IAF peak of 10 Hz would have a bandwidth ranging from 8.33 Hz to 11.67 Hz. Power was extracted from two symmetric regions of interest precisely in PR (P6, P8, PO4, O2) and PL locations (P5, P7, PO3, O1) in order to replicate as closely as possible the original EOI electrodes used in Thut et al., (2006). Power imbalance was computed according to the formula:

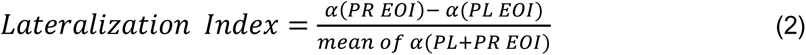

where α (PL EOI) and α (PR EOI) are the average of α-power over left and right electrodes of interest, respectively. Equation (2) leads to smaller (negative) values where α-activity is more prominent over the left hemisphere than the right (α (PL EOI) > α (PR EOI)) and to larger (positive) values for the opposite pattern (α (PL EOI) < α (PR EOI)). According to theory and previous findings, values of LI reflecting attention directed to the right hemifield should be larger than LI values reflecting leftward directed attention.

Finally, we also checked whether there was any relationship between the α-power imbalance and the contra-ipsi difference of PLV for each attended location. We explored the correlations between α-lateralisation indexes and the effect in PLV contra-ipsi differences at the pre-target (-200 to 0 ms) and post-target (200 to 400 ms) windows using Pearson correlations.

### Statistical analyses

A one-tailed nonparametric Monte Carlo permutation test was computed to determine significant differences in PLV between networks for each attended location (Mostame et al. 2019). For each participant, the *attended right* or *left* labels were randomly assigned to trials, and surrogate PLVs were calculated from the resulting dataset. This process was repeated 10,000 times (iterations) to create a null distribution of PLV values. The obtained p-value corresponded to the proportion of surrogate iterations with a contra-ipsi difference larger than the actual measured value (one-tailed test). This process was performed on every time window defined in the previous section. For the group analysis, the procedure was equivalent, but surrogate PLV distributions were averaged across participants before the statistical test.

For the statistical assessment of the α-power imbalance over time between *attended left* and *attended right* trials, we performed a cluster-based permutation test procedure (100,000 randomisations) for each participant and at the group-level (one-tailed permutation test) (Maris and Oostenveld 2007; Meyer et al. 2021). We assessed that lateralisation indexes for *attended right* and *attended left* trials were two significantly different distributions by applying a one-tailed t-test (independent samples) with α-level = 0.05 for each participant. At group-level, we performed a one-tailed paired t-test with the mean lateralisation indexes for *attended right* and *attended left* trials for each participant with α-level = 0.05. Correlations between α-power imbalance and the contra-ipsi difference of PLV were corrected for multiple comparisons by applying the False Discovery Rate (FDR) of Benjamini and Hochberg (Benjamini and Hochberg 1995).

## RESULTS

### Behavioural results

Five participants who presented equivalent detection and discrimination rates for stimuli appearing at cued and un-cued locations were discarded from the analysis, leaving a total of 10 participants. As expected, behavioural results showed that the detection rate calculated based on both the detection response (Yes/No) and the discrimination response (Left/Right; chance level at 0.25) was superior for cued (attended) trials 0.68 ± SEM=0.02 compared to un-cued (unattended) ones 0.46 ± 0.04 (see, **Figure 1B**). The pattern on each hemifield was equivalent: on the left hemifield attended = 0.68 ± 0.03 and unattended = 0.47 ± 0.03; for the right hemifield attended = 0.67 ± 0.03, and unattended = 0.44 ± 0.06. We used one tailed t-tests to assess that performance was above chance level (25%) for each of the conditions (attended and unattended) and hemifields separately: Attended Left trials (0.68±0.11, p-value=2·10^-7^, t(9)=12.593), Attended Right trials (0.67±0.11, p-value=3·10^-7^, t(9)=12.226), Unattended Left trials (0.47±0.08, p-value=5·10^-6^, t(9)=8.876) and Unattended Right trials (0.44±0.20, p-value=0.06, t(9)=3.117).

### Target-locked long-range α synchrony

Here, we describe the results from the target-locked analysis, carried out to reproduce Sauseng et al.’s (2005) findings. Long-range synchrony was estimated using PLV between frontal EOI and each of two lateralised parietal EOI. **Figure 2** shows the group-level connectivity analysis of the upper α-band (9.54 - 14.31 Hz). Phase coupling is depicted as the mean across the pre-target window (-200 to 0 s) and the post-target window (200 to 400 ms), as well the temporal course (from to -500 to 500 ms). Regarding the left fronto-parietal network (**Figure 2A** **left**), PLV was consistently higher when attention was directed rightward (contralateral) than leftward (ipsilateral) in both pre-target and post-target windows, although the PLV difference only reached significance in the post-target window (*p* < 0.05). Regarding the right network (**Figure 2A** **right**), PLV was stronger when attention was directed leftward (contralateral) than rightward (ipsilateral) in the post-target window, whereas the pre-target window does not show this difference. Neither window, however, emerged as significant. This pattern generally replicates Sauseng et al. (2005) results, as indicated by the dashed lines in **Figure 2A** representing the mean phase-coupling from their study. Lower panels in **Figure 2A** display the temporal course of phase coupling to provide a time-resolved illustration of the phase-coupling effect. For the *attended right* condition, PLV values in the left network should be higher than PLV values for the *attended left*. The inverse pattern should hold in the right network. Moreover, **Figure 2B** presents the PLV with side of attention collapsed as contra- and ipsilateral with respect to the corresponding network. Individual PLV values, marked as black dotted line s, exhibit a consistent contra- to ipsilateral increase in the post-target window. Group-level statistical analysis further showcased a significant difference limited to this time window (200 to 400 ms, *p <* 0.05). This result was controlled by avoiding the pre-processing band-pass filter which may affect phase estimation, and by computing a Hjorth filter to avoid the effects of volume conduction (Hjorth 1975). Both analyses maintained the significant differences between contralateral and ipsilateral PLV (*p* < 0.05).

**Figure 2.**
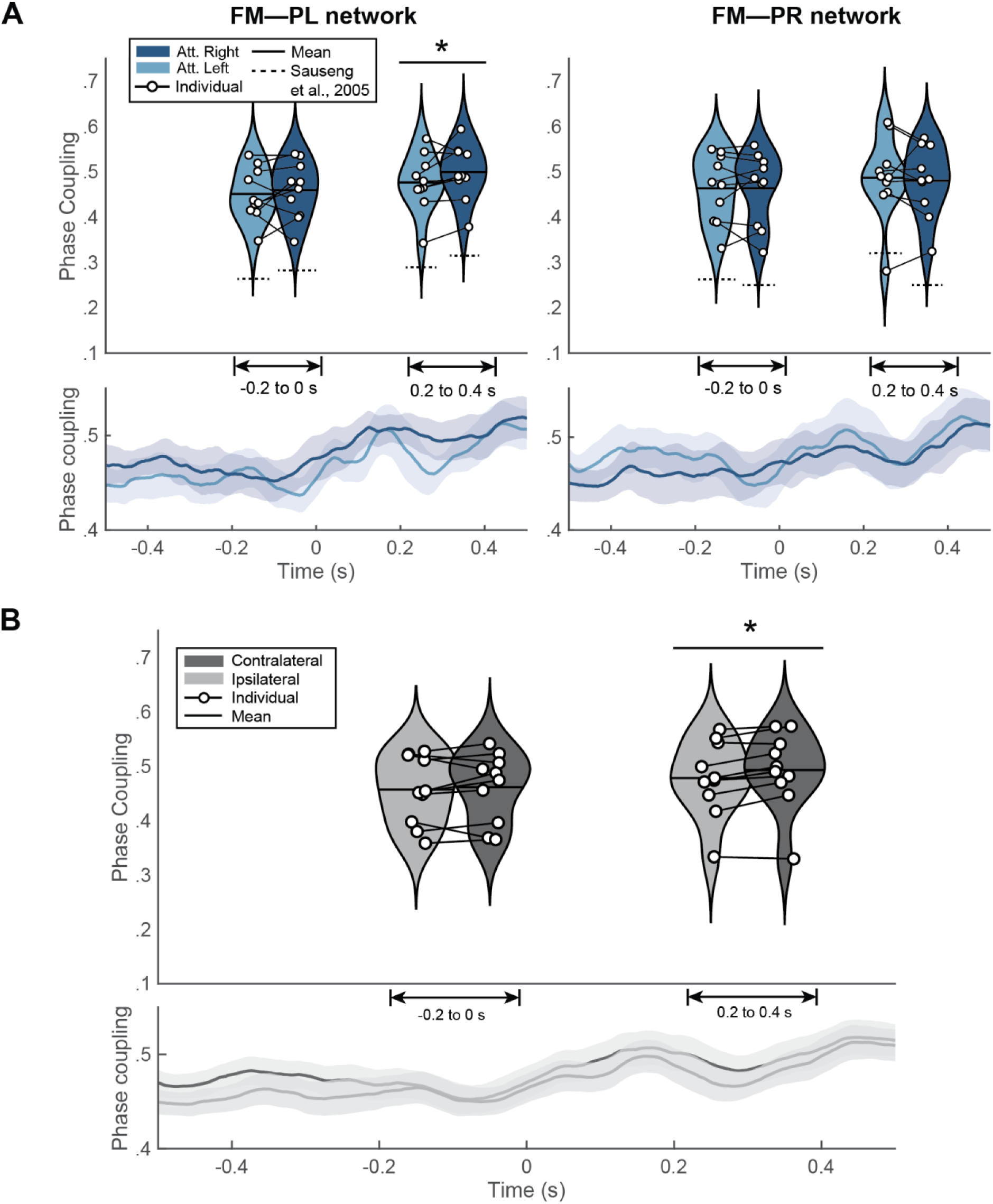
Target-locked results. **(A) Target-locked results of the phase-coupling for *attended left* (light blue) and *attended right* (dark blue) in FM-PL and FM-PR networks.** The lower panels depict the cross-trial average time course (± shaded SEM) of PLV in both conditions (*attended left* and *attended right*). Upper panels present the binned violin plots (mean and median) of the pre-target window (-200 to 0 ms) and the post-target window (200 to 400 ms); \**p* < 0.05. **(B) Target-locked results collapsed as either ipsilateral (FM-PL network and *attended left*; FM-PR and *attended right*) or contralateral (FM-PR network and *attended left*; FM-PL and *attended right*).** The lower panel shows the cross-trial average time course (± shaded SEM) of PLV in ipsilateral (light grey) and contralateral (dark grey) conditions. The upper panel exhibits the distribution of individual PLV with a violin plot, superimposed by the mean and the contra- to ipsilateral differences between individual PLV; \**p* < 0.05.

At individual level, only 3 out of 10 participants showed significant contralateral PLV increase (P02, *p <* 0.01; P05, *p <* 0.01; P07, *p <* 0.01; see **Figure 2-1**). The lack of a significant group-level effects in the pre-target window is consistent with individual phase coupling, as a multiple subject present a trend in the opposite direction as expected (i.e., ipsilateral over contralateral PLV; see **Figure 2-1**). We further assessed single-subject synchronization through the phase linearity measurement (PLM) as it has been recently reported to be a robust metric for trial-level connectivity (Baselice et al., 2018). We did not find any significant effects in any participant (*p* > 0.05; see **Figure 2-2**).

### Cue-locked long-range α synchrony

In the previous section, we replicated the results as in Sauseng et al., (2005). The findings from here onwards correspond to original results to ascertain whether attention-based long-range connectivity during the attention-orienting period could be a reliable signal for BCI control. We explored the cue-to-target interval before target presentation (500 ms to 1500 ms after cue onset). Considering that the cue indicates the hemifield to which participants should voluntarily lateralise attention, differences in contralateral and ipsilateral connectivity may potentially emerge in this time window. So far, we have seen that attention shifts had significant consequences on behaviour and target processing (post-target connectivity). At the group level, however, no significant difference between contralateral and ipsilateral connectivity in the upper α-band was found in any of the five 200 ms time windows considered in the cue-to-target period (see **Figure 3A**). At the individual level, 7 participants had a significant contralateral PLV increase in at least in one window (see **Figure 3-1**). However, only one participant (P10) showed this effect in all time windows and, furthermore, did not present a significantly higher contralateral connectivity in pre-target and post-target time windows of the target-locked analysis.

**Figure 3.**
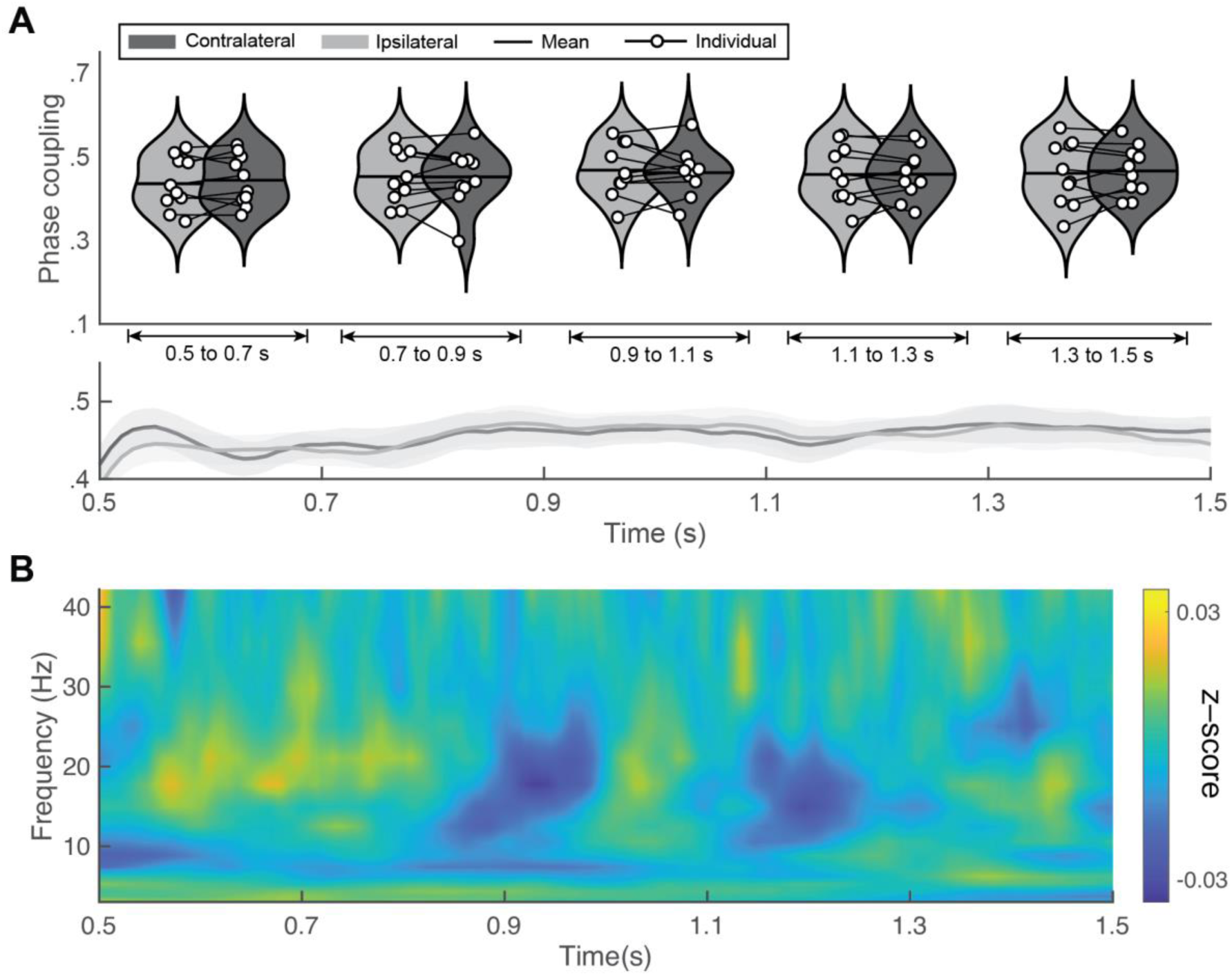
Cue-locked results. **(A) Group-level results of upper-alpha PLV.** Upper panel shows phase coupling for ipsilateral (light grey) and contralateral (dark grey) sides in time-windows of 200 ms from the cue-locked interval (500 ms to 1500 ms after cue presentation). Lower panel shows mean and standard error of the mean (SEM) of the PLV values. **(B) Exploratory analysis of PLV differences.** Group-level temporal evolution of the z-scored difference between contralateral and ipsilateral PLV for each frequency band (2.4 - 42 Hz with 16 logarithmic steps). Z-score values range from - 0.03 to 0.03.

We chose the upper α-band a priori given Sauseng et al. (2005)’ findings, as well as the effects in the target-locked analyses from the present dataset. However, we conducted additional analyses to explore other frequencies (between 2.4 and 42 Hz) in search of differences between contralateral and ipsilateral PLV (see, **Figure 3B**). Values were collapsed as the difference between both measures (contra-ipsi) and z-scored. Over time, neither clear trends across frequencies nor apparent increases were observed in contralateral or ipsilateral connectivity. Individual results showed the same trend and did not present relevant PLV patterns in any participant beyond those from upper α-band findings in P10 (see **Figure 3-2**).

### Classification

The results are hardly promising in generalising the use of long-range connectivity for BCI control. However, BCI protocols are often very sensitive to individual patterns. Here, we intended to seek a proof-of-concept, from at least a single participant. With this goal in mind, we attempted single-trial classification, as either *attended right* or *attended left*, according to cue-locked connectivity patterns. We selected the participant (P10) for whom we found significant connectivity differences in the cue-to-target time window of the cue-locked analysis. The total number of trials was 338.

We carried out a validation of cross-time PLV in the target-locked window to understand whether this metric could replicate group-level differences between contra- and ipsilateral networks found through cross-trial PLV. These results can be seen in **Figure 4A**. Statistical analysis showed no significant differences between contra- and ipsilateral scenarios in either time window. Individual values were also non-significant (see **Figure 4-1**). Considering the large parametric landscape of SVM implementations, we optimised the gamma and margin parameters of a Gaussian kernel (see **Figure 4B**). From a qualitative perspective, no clear maximum validation accuracy values emerge from the landscape, although quantitative analysis identified minimum values of margin and γ to be used on the test set in every fold. The lack of a clear minimum suggests that the model may be unable to classify individual trials regardless of the parametric values.

**Figure 4.**
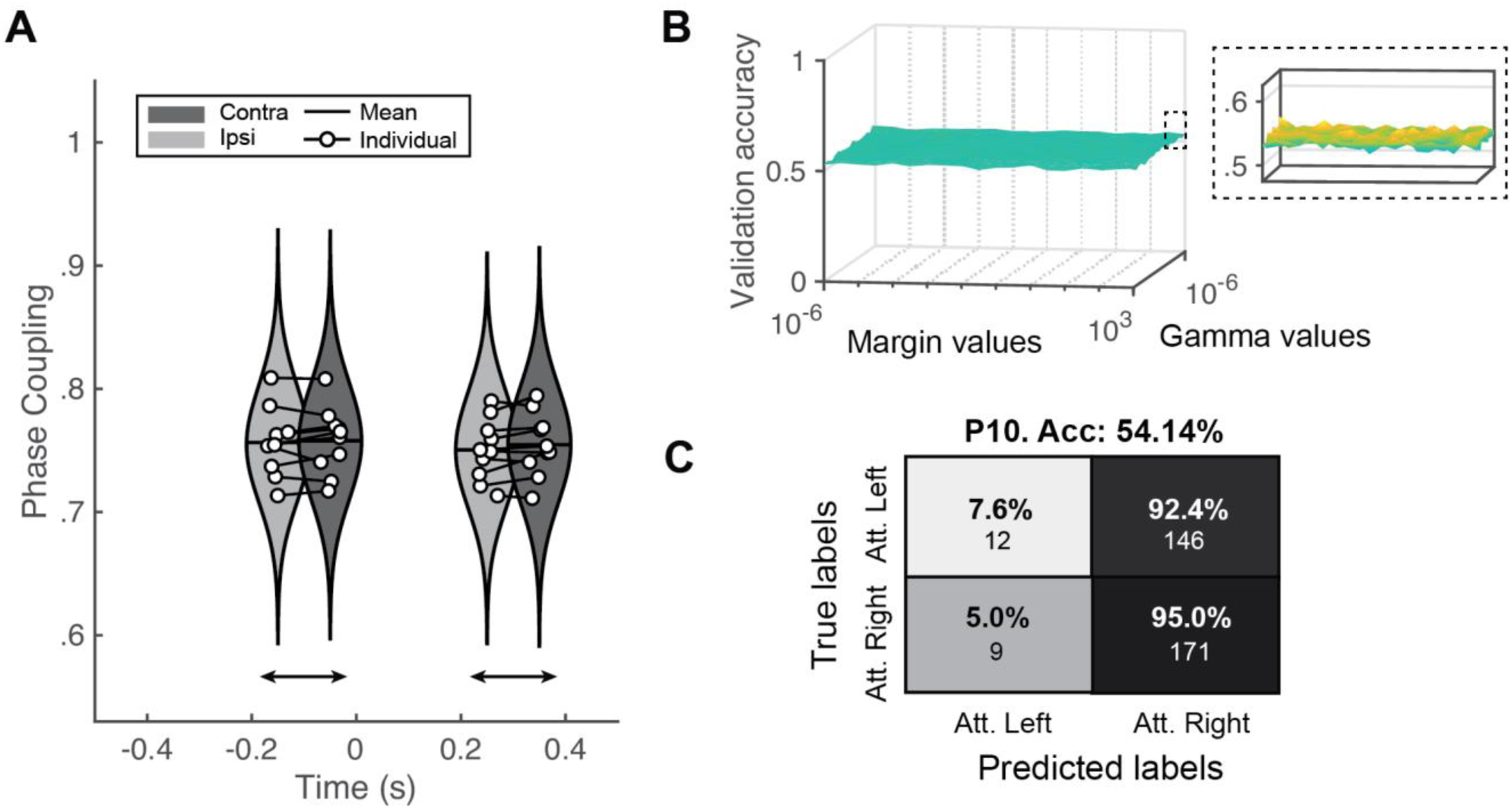
Classification outcomes. **(A) Cross-time PLV reality check.** Replication of results from Fig. 2 calculating PLV across time points rather than across trials. **(B) Optimisation results of gamma and margin parameters of the Gaussian kernel SVM.** Ten-fold validation accuracies with varying margin values (x-axis) and gamma values (y-axis). Inset shows a detailed view of the z-axis. **(C) Confusion matrix of the classification outcomes for one participant.** Y-axis represents ground truth labels (*attended right* or *attended left*) and x-axis represents the classifier’s outcomes. Percentages represent the fraction of correctly classified trials of each condition (i.e., each row sums to 100%). Under the percentage is the gross number of classified trials.

Ten-fold cross-validation was carried out to maximise the available data and improve the classification accuracy. Single trials predicted as either *attended right* or *attended left* were contrasted with the actual cue direction in each trial. Classification outcomes are shown in **Figure 4C**, which resulted in virtually chance-level sorting (0.541). The confusion matrix displays the distribution of each class, revealing the skewed distribution of values towards *attended right* labels, which is far from the ideal clustering along the diagonal of the matrix. Finally, we employed two additional algorithms to classify both *attended right* and *attended left* trials. These consisted of shrinkage linear discriminant analysis (sLDA) and Riemannian minimum distance to the mean (RMDM), as they are shown to work well in small training sets (Lotte et al., 2018). Both decoding techniques yielded chance-level results (see **Figure 4-2**)

### Inter-hemispheric power imbalance

As a reality check on the dataset, we addressed whether there was a difference in the α-power inter-hemispheric imbalance between *attended left* and *attended right* trials. We performed the cue-locked analysis at the group level, using the Lateralization Index (LI) described by Thut et al. (2006) (see **Figure 5A**). On average, the lateralisation index was significantly different between *attended right* and *attended left* in the expected direction (*p <* 0.01, Cohen’s d = -0.8356). At the individual level, 7 out of the 10 participants showed a significant difference in lateralisation index between the two attention conditions (*p <* 0.05; see **Figure 5-1**). We also performed a time-resolved version of this analysis within the cue-to-target window. A cluster-based permutation test (**Figure 5B**) showed significance within two time periods, from 0.66 to 0.82 s and 1.34 to 1.5 s. At the individual level, only for one participant (P01), the cluster-based permutation test revealed a significant cluster over time from 0.6 to 1 s (see **Figure 5-1**). These results are consistent with the results of previous studies (e.g., Tonin et al. 2012; Thut et al. 2006), at least at the group level. It is more challenging to compare single-subject data with other studies, as it usually is not reported or statistically analysed.

**Figure 5.**
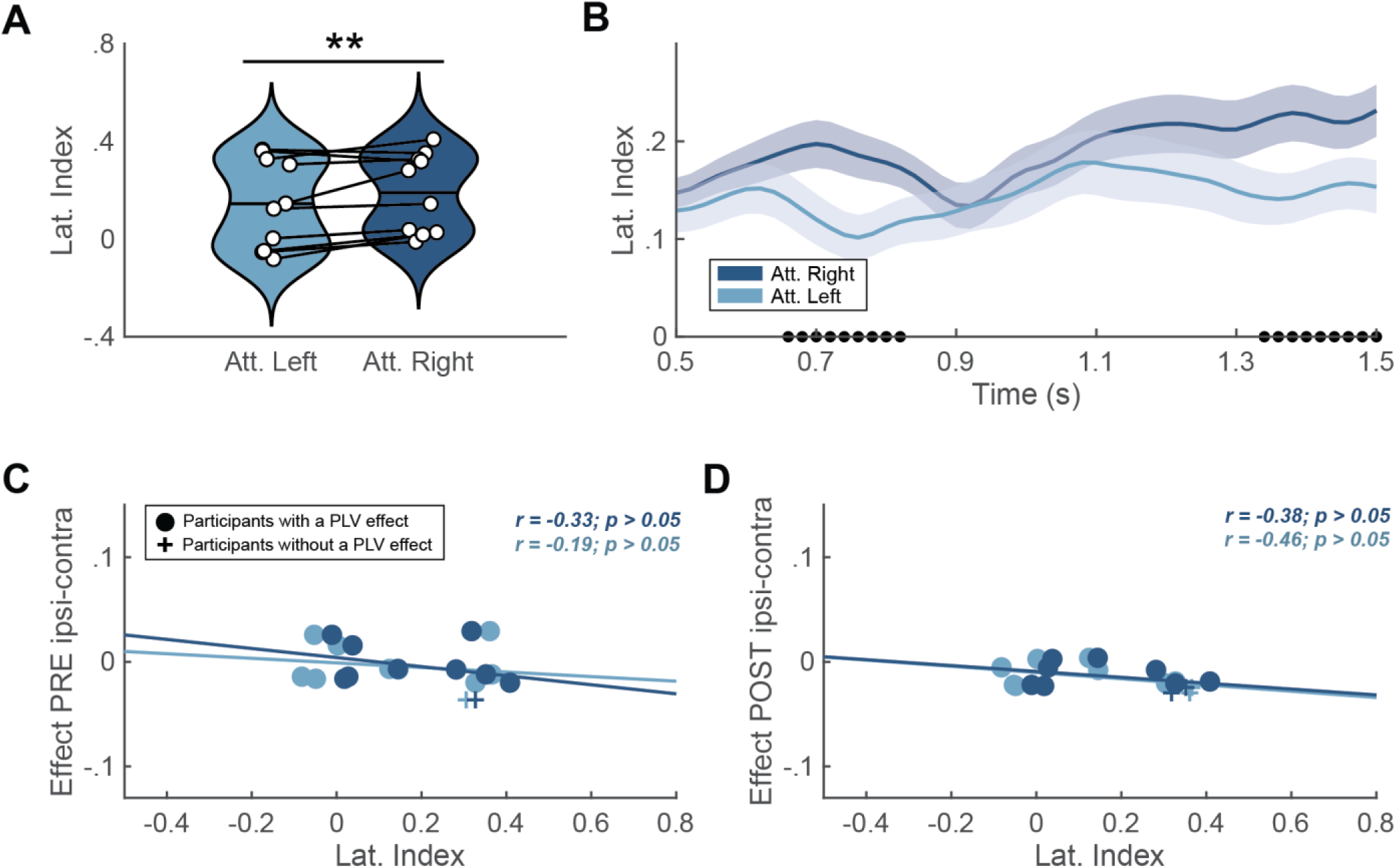
Lateralisation index reality check. **(A) Averaged lateralization index for *attended left* (light blue) and *attended right* (dark blue; \**p* < 0.05; **p < 0.01).** White dots denote individual scores, and horizontal line indicates the group mean. **(B) Lateralisation index (mean ± SEM) over time.** Solid lines and shaded areas represent mean and standard error of the mean (SEM) interval, respectively. Dots on in the x-axis denote the significant difference over time between *attended left* (light blue) and *attended right* (dark blue) via cluster-based permutation test. **(C-D) Lateralisation indexes and the difference of contra- to ipsilateral PLV for *attended left* (light blue) and *attended right* (dark blue) at the pre-target window (C) and the post-target window (D).** At the pre-target the correlations for *attended right* (*r* = -0.33, *p* > 0.05) and *attended left (r* = -0.19, *p* > 0.05) did not reach significance and neither did the correlations for *attended right* (*r* = -0.38, *p* > 0.05) and *attended left r* = -0.46, *p* > 0.05) at the post-target window. Crosses denote participants with a significant effect in PLV contra-ipsi differences at the pre-target window (-200 to 0 ms; P05) and the post-target window (200 to 400 ms; P02 and P07). Dots represent the rest of the participants.

Finally, we explored the potential correlation between α-power inter-hemispheric imbalance measured with the lateralization index and α-phase coupling for each attended location (see **Figure 5 C-D**). In the pre-target window (**Figure 5C**), the correlations for *attended right* (*r* = -0.25, *p* > 0.05) and *attended left* (*r* = -0.13, *p* > 0.05) did not reach significance. Neither did the correlations for *attended right* (*r* = - 0.44, *p* > 0.05) and *attended left* (*r* = -0.42, *p* > 0.05) at the post-target (**Figure 5D**) window. A visual inspection indicated that participants showing an effect in PLV contra-ipsi differences are below the correlation fit in pre-target and post-target windows, suggesting that those participants have a more negative effect in PLV contra-ipsi differences.

## DISCUSSION

The present study addressed the relationship between shifts in visuospatial attention and the lateralisation of α-band coherence between frontal and parietal electrodes, to assess their feasibility as a control signal in BCI. Previous studies, using group-averaged multi-trial analyses, found increased long-range α-synchronisation in the hemisphere contralateral to the attended hemifield, and suggested that it reflects top-down mechanisms of visual spatial attention (Sauseng et al., 2005; Doesburg et al., 2009). We reasoned that if contra- to ipsilateral differences in synchronisation would emerge as a result of endogenous top-down mechanisms, they should be present following cue presentation as participants shift their attention. This hypothesis stems from how instructing participants to shift their attention laterally before target appearance engages frontoparietal visual processing pathways (Corbetta and Shulman 2002; Hopfinger et al. 2000; Asplund et al. 2010). Here, we sought proof that long-range neural synchronisation engaged in this network could be used for BCI control on a trial-by-trial basis.

In attention-orienting protocols, the cue-to-target period offers the possibility of implementing a BCI control in anticipation of the target appearance. This would open the possibility of designing active BCI systems controlled by the user’s voluntary decision to attend left/ rightward covertly. Therefore, our study employed long-range α-synchronisation in the frontoparietal network (FPN) as means to investigate whether this brain measure could potentially discriminate attended locations of the left/right visual field.

We found significant group-level differences in contra- to ipsilateral long-range α-synchronisation around target onset, replicating Sauseng et al. (2005). These results demonstrate the involvement of lateralised long-range α-synchrony along the FPN during the post-target period and especially reveal the potential of EEG to grasp these effects, at the group level. However, similar differences in fronto-parietal synchrony were not observed during the cue-to-target time window, which was the time of interest for BCI purposes. We also extended the cue-locked analysis to other frequencies outside the α-band, with equally negative results. Finally, given the high individual variability of single-trial analysis outcomes, we attempted to classify the individual trials of one selected participant for whom significant synchronisation differences following cue presentation were found, as a benchmarking process. The results nevertheless rendered chance-level classification. Below, we discuss how these results may be influenced by various methodological aspects (e.g., different time windows, classifier’s input metric) and how they fit into state-of-the-art literature. Please note that because the focus of our study was on single-trial analysis, the sample size was relatively small for the group analyses (n = 10). Although this sample size was sufficient to confirm previous findings on long-range α-synchronisation and lateralization index (Sauseng et al., 2005; Thut et al., 2006), the negative results of the group analyses should be interpreted with caution.

### Fronto-parietal network synchronisation characterises visuospatial attention

A result from our study is that long-range α-synchronisation within the FPN was associated with the consequences of visuospatial attention orienting, in line with its putative role in this cognitive process (Jensen et al. 2015; Sacchet et al. 2015; Doesburg et al. 2009; Siegel et al. 2008). We observed significant increase in contralateral vs. ipsilateral upper α coherence for targets appearing at the attended location. According to the current attention theories, the mechanism underlying this finding may be inherently related to top-down processing. More specifically, frontal regions such as the frontal eye fields (FEF) and the intraparietal sulcus (IPS) may modulate attention by causing a state of α-band desynchronisation in the visual cortex contralateral to attended hemifield (Corbetta and Shulman 2002; Kastner and Ungerleider 2000; Helfrich et al. 2018; Capotosto et al. 2009; Marshall et al. 2015). This explanation further aligns with the well-established evidence that contralateral α-power suppression (also reproduced in our results) enables visual stimuli processing in the attended location (Doesburg et al. 2009; Thut et al. 2006; Yamagishi et al. 2003; Babiloni et al. 2006; Foxe and Snyder 2011; Klimesch et al. 2007; Lange et al. 2013), and that cyclic phase-dependent inhibition in low-level visual cortex dictates behavioural performance (i.e., reaction times) (Haegens et al. 2011; Klimesch 2012; Jensen et al. 2014; Samaha et al. 2015; VanRullen 2016). Both accounts fit with the idea that local α-power and long-range α-synchronisation may have separate roles in attention and perception (Bonnefond et al. 2017; Palva and Palva 2007, 2011; Sadaghiani and Kleinschmidt 2016).

Our results of the increased contralateral synchronisation within the FPN replicate the work of Sauseng et al. (2005) and validate our methodology and analysis pipeline (e.g., time-frequency analysis, synchronisation metric), setting the ground for the intended proof of concept test regarding transference to BCI. However, lateralised fronto-parietal connectivity patterns in attentional and perceptual disposition remain challenged in the literature together with the role of α power/phase (Ruzzoli, Torralba et al. 2019; van Diepen et al. 2019; Antonov et al. 2020; Keitel et al., 2022). Lobier et al. (2018) found that α-synchronisation was associated with visuospatial attention but revealed distinct lateralisation patterns regarding the visual system and top-down attentional networks. They showed stronger ipsilateral synchronisation within the visual system (in line with Siegel et al. 2008; Doesburg et al. 2009) but no consistent lateralisation in long-range networks, suggesting their different involvement in visuospatial attention. A study by D’Andrea et al. (2019) found a modulation of frontoparietal α-β cross-frequency synchronisation during attention orienting, but not in α-synchronisation alone. Further, this cross-frequency connectivity pattern was strongly associated with right hemisphere frontal dominance, in line with Heilman and van den Abell (1980) and Zago et al. (2017). This finding agrees with previous evidence of the crucial role of the right FEF in top-down attentional modulation (Esterman et al. 2015; Hung et al. 2011; Silvanto et al. 2006; Veniero et al., 2021), supported by evidence using TMS (e.g., Capotosto et al. 2009). In light of this evidence and our results, the exact relationship between contralateral frontoparietal α-synchronisation and shifts in attention orienting is still unclear. Positive findings, however, such as the ones in the present study using a target-locked analysis, represent a basis for exploring earlier time windows capable of shedding light on the mechanism underlying FPN α-synchronisation.

Correlations between long-range α-synchronisation and individual reaction times in visuospatial tasks suggest this neural correlate may be observable at a single-subject level (Lobier et al., 2018). However, significant group-level target-locked dynamics of increased synchrony did not transfer to all individuals in our study. The observed variability may be partially explained by individual anatomical differences in the neural substrate of attention (e.g., superior longitudinal fasciculus) (Marshall et al., 2015). Findings employing magnetic resonance imaging (MRI) suggest that volumetric differences in these structures impact local visual cortex oscillations, leading to variability in EEG traces (Marshall et al. 2015; D’Andrea et al. 2019). However, this variability of individual results is challenging to set in the perspective of previous research simply because published studies do not report single-subject statistics. Ultimately, the outcomes of this study leave an incomplete understanding of whether there is a reliable group effect that does not extend to all individuals or, contrarily, whether individual effects of specific participants are large enough to induce a group-level finding in previous research.

### Lateralized patterns of α-synchronisation appear in target-locked but not cue-locked analysis

In our study, long-range α-synchronisation presented contralateral increases at the post-target (200 to 400 ms, with t = 0 as target appearance) and the pre-target window (-200 to 0 ms), but only the former time window resulted significantly. This result is slightly different from Sauseng et al. (2005), who observed significant increases in contralateral synchronisation within the FPN network at both time windows. However, the numerical differences were in the same direction in both studies, leaving the possibility that statistical significance be just due to a lack of statistical power. Another potential explanation for the absence of significant findings at the pre-target window may be the difference in experimental paradigms. The task employed here had a longer post-cue interval ranging from 2000 to 2500 ms (jittered between trials), compared to Sauseng et al. (2005) (i.e., 600-800 ms). If participants shifted attention at varying times from cue onset up to target appearance, this might explain why we could not capture the effect in anticipatory visuospatial attention.

In cueing paradigms, bottom-up integration of cue information through sensory pathways precedes top-down modulation of visuospatial attention (Simpson et al., 2011). The temporal course of voluntary directed attention is thought to begin only after 150 ms from cue onset and involves frontal regions approximately after 350 ms. Furthermore, from 400-500 ms onwards, frontal and parietal regions are thought to be involved in attentional shifting and target discrimination (Simpson et al. 2011). Thus, if the FPN does present direction-specific synchronisation, we anticipated this would appear from about 500 ms after cue onset onwards. Contrary to what we expected, we did not observe any significant contra- to ipsilateral differences in the cue-to-target time windows (500 to 1500 ms after cue onset). Previous studies employing a similar time window showed lateralisation patterns in parietal regions in α and β bands (Siegel et al. 2008; Pantazis et al. 2009) and frontoparietal lateralisation in low and high-frequency bands (Green and McDonald 2008; Gregoriou et al. 2009). Therefore, we extended our cue-locked analysis to other frequencies but again obtained no significant contra- to ipsilateral differences. Note that PLV values were averaged across 200 ms windows, and this excludes, to a certain extent, the confound of frontal and parietal regions having different activation over time. Altogether, despite the evidence across multiple frequencies of synchronisation in the cue-to-target time window, we did not find patterns of lateralised cue-locked connectivity within or outside the α-band.

Our negative results in the cue-locked analysis may align with the notion that late periods after cue onset are associated with direction-specific activity in parieto-occipital regions but not in frontal regions (e.g., FEF) (Doesburg et al. 2009; Simpson et al. 2011). Long-range α-synchronisation may, therefore, be associated to an initial shift of attention (shortly after cue presentation) and later (close to target presentation) to attention maintenance at the directed hemifield (Lobier et al. 2018; Kastner and Ungerleider 2000; Hopfinger et al. 2000; Grent-’t-Jong and Woldorff 2007). This idea resonates with the essential question formerly posed by Sauseng et al. (2005), debating whether frontal involvement in long-range α-synchronisation is a causative or consequential correlate of posterior activation. Furthermore, it motivated the exploration of cue-locked intervals where bottom-up and top-down processing may have elicited stronger effects on α-band synchronisation.

Finally, to ensure participants correctly lateralised their attention during the cue-to-target interval, we carried out a reality check by calculating the α-power imbalance using the lateralisation index during this period (Thut et al. 2006). There was a clear difference in the averaged lateralisation index during the time course between 500 and 1500 ms at group-level. We further employed the lateralisation index to perform an exploratory analysis of its relationship with the difference in α-synchronisation between contra- and ipsilateral networks. Considering lateralised local α activity and lateralised long-range α-synchronisation are both relevant in successful attention orienting, we explored whether these two mechanisms would have had a significant positive correlation. Therefore, individuals with high lateralisation index values should also present lateralised synchronisation within the FPN. In contrast to our expectations, there was no significant correlation between these two metrics, neither at the pre-target nor the post-target time windows.

Ultimately, we did not observe a significant increase in contralateral long-range α-synchronisation in the five 200 ms bins following cue onset. This time frame offered potential as it occurs much before target appearance and could be robustly employed in a covert visuospatial BCI decoder. By expanding our analysis to several frequencies and carrying out the aforementioned reality checks, we conclude that PLV measured from EEG may not serve as a reliable metric in capturing direction-specific synchronisation from frontal to posterior regions, despite this evidence being present in parietal to occipital synchrony (Doesburg et al. 2009).

### EEG estimates of long-range α-synchronisation may not serve as a reliable control signal for BCI

The use of long-range α-synchronisation to decode attentional direction yielded chance-level results. We employed 200 ms time bins of contralateral and ipsilateral FPN connectivity as input in an SVM classifier. Non-linear SVMs are widely employed in decoding cognitive neural correlates of behavioural states (Lotte et al., 2007). Furthermore, SVMs outperform other classifiers, such as artificial neural networks, non-linear Bayesian estimators, and recurrent reservoir networks (Astrand et al. 2014a). We also employed sLDA and RMDM classifiers, as they have low computational cost, require small training sets, and perform well in real-time applications (Lotte et al., 2018), with no success.

Prior work using SVMs, mainly centred around primate models and invasive recordings, successfully decoded the attentional spotlight from frontal sites (Gaillard et al. 2020; Tremblay et al. 2015; Esghaei and Daliri 2014). Clearly, these methods (i.e., LFP, intracranial-EEG) have a higher signal-to-noise ratio (SNR) compared to non-invasive imaging. However, the objective of the present study was to offer a BCI proof of concept using α-synchronisation as a control signal. Therefore, a non-invasive and portable technique must be employed. Other non-invasive modalities such as functional magnetic resonance imaging (fMRI), where the temporal resolution is too low for real-time implementations, or magnetoencephalography (MEG), where the equipment is expensive and requires a magnetically shielded room (as fMRI), have limited potential transfer in out-of-lab applications. Contrarily, EEG is an affordable imaging modality with a straightforward setup which provides high temporal resolution and portability. However, the inconvenience of using EEG is a low spatial resolution and a low SNR. Despite this, decoders have been commonly employed in EEG-BCI design employing parieto-occipital power changes in α-band activity to predict covert visuospatial attention tasks (Tonin et al. 2013; Treder et al. 2011; van Gerven et al., 2009). The integrated approach between frontal and parieto-occipital attentional decoding based on α-synchronisation, however, has not been attempted. Here, we found that cue-locked synchronisation enclosed in the FPN α-band is insufficient to determine the attentional location at EEG single trial level. This may be due to an inherent lack of connectivity in the cue-to-target interval, or else more likely, the poor sensitivity of the EEG to register synchronisation patterns.

Another potential reason to explain the failed classification of cue-locked FPN connectivity at single-trial level may be the change in PLV calculation from trial-average to single-trial. Standard cognitive research employs multiple trials to estimate consistent findings on electrophysiological markers (M/EEG). Instead, BCIs need robust and accurate estimates in a single-trial fashion and thus require a trade-off between spatial (i.e., single-channel decoding is preferred) and temporal resolution. PLV is a measure of consistency across multiple trials and cannot serve as a single-trial control signal. Therefore, we computed PLV across time points within the same trial. This new measure is also referred to in the literature as the inter-site phase clustering (ISPC) and may represent a different underlying process than that captured by classic PLV (Cohen 2015). This prompts the question of whether long-range α-synchronisation is incapable of decoding the attended location, or rather the single-trial nature of IPSC over time is responsible for this.

In sum, long-range α-synchronisation within the FPN estimated with EEG may not serve as a control signal for BCI. This limitation may be due to incomplete information on neural correlates due to the lack of cross-frequency analysis or the computational techniques surrounding ISPC over time.

## CONCLUSION

We found direction-specific contralateral patterns of upper α-synchronisation (i.e., PLV) within the FPN following target appearance in a covert visuospatial task. This finding, however, did not extend to pre-target or cue-to-target time windows. The modulatory role of α-synchronisation in anticipatory attention through frontal, parietal and occipital regions suggests that PLV may not constitute a reliable metric for this top-down visual processing. Furthermore, chance-level classification resulting from using this metric in an SVM indicates that long-range α-synchronisation computed with EEG may not be a suitable control signal for BCI.

---

**Figure 2-1.**
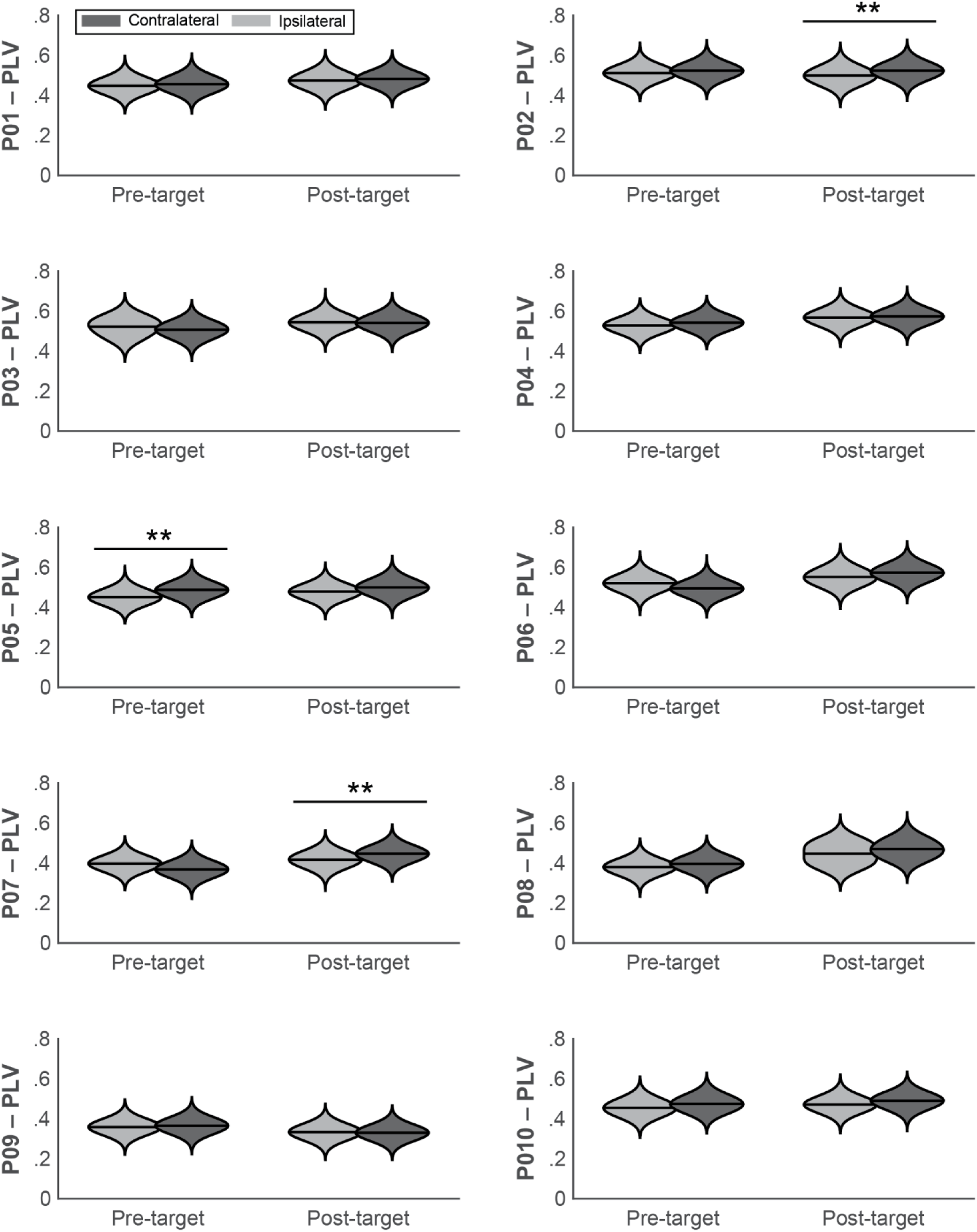
Individual results of target-locked PLV index. Violin plots represent the phase locking values (PLV) averaged over the pre-target (-200 to 0 ms, t = 0 as target appearance) and post-target time window (200 to 400 ms). Ipsilateral (FM-PL network and *attended left*; FM-PR and *attended right*) or contralateral (FM-PR network and *attended left*; FM-PL and *attended right*) scenarios are exhibited as either light grey or dark grey, respectively. \**p* < 0.05, \*\**p* < 0.01, \*\*\**p* < 0.001.

**Figure 2-2.**
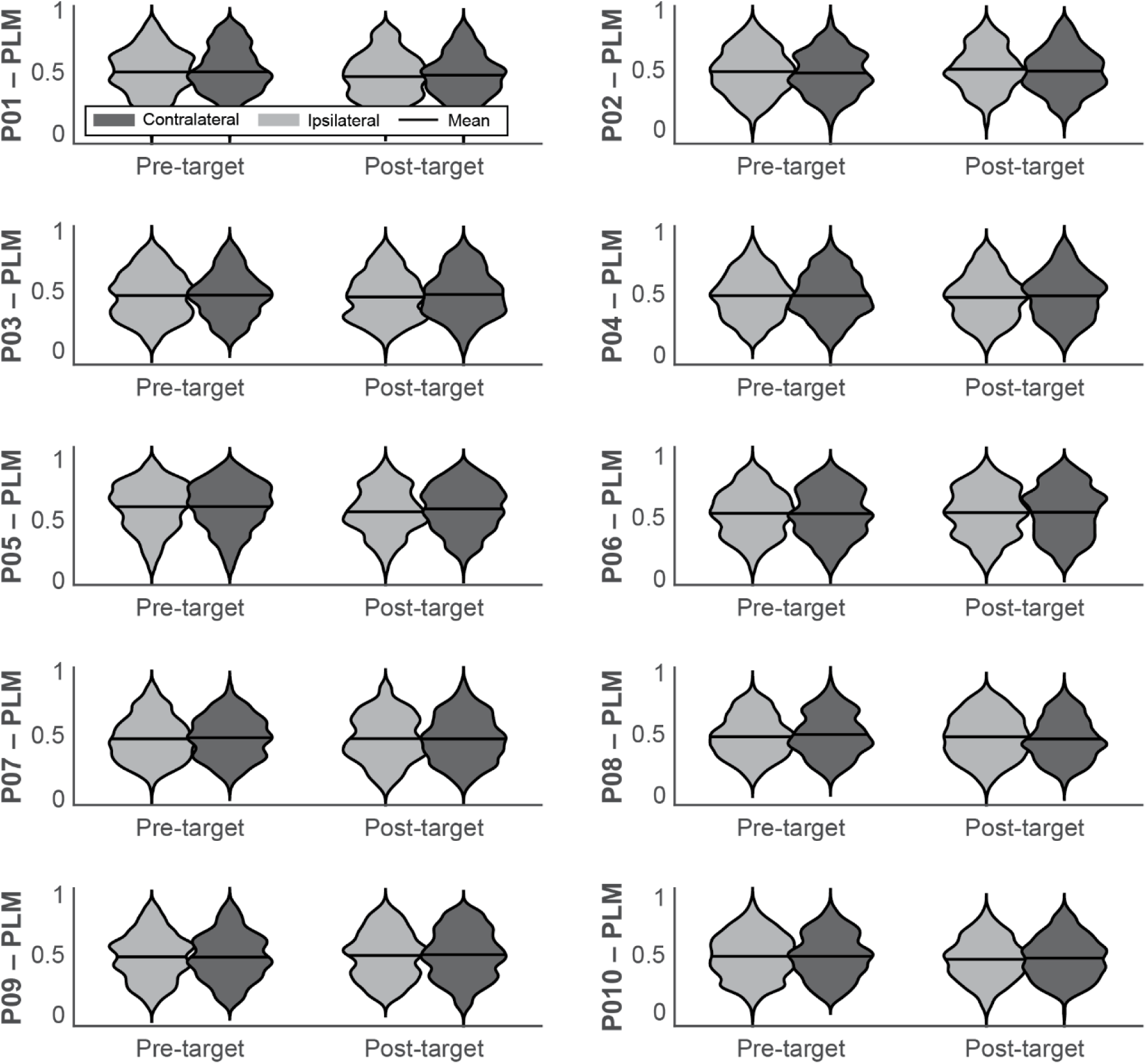
Individual results of target-locked PLM index. Violin plots represent the phase linearity measurement (PLM) over the pre-target (-200 to 0 ms, t = 0 as target appearance) and post-target time window (200 to 400 ms). Ipsilateral (FM-PL network and *attended left*; FM-PR and *attended right*) or contralateral (FM-PR network and *attended left*; FM-PL and *attended right*) scenarios are exhibited as either light grey or dark grey, respectively. \**p* < 0.05, \*\**p* < 0.01, \*\*\**p* < 0.001.

**Figure 3-1.**
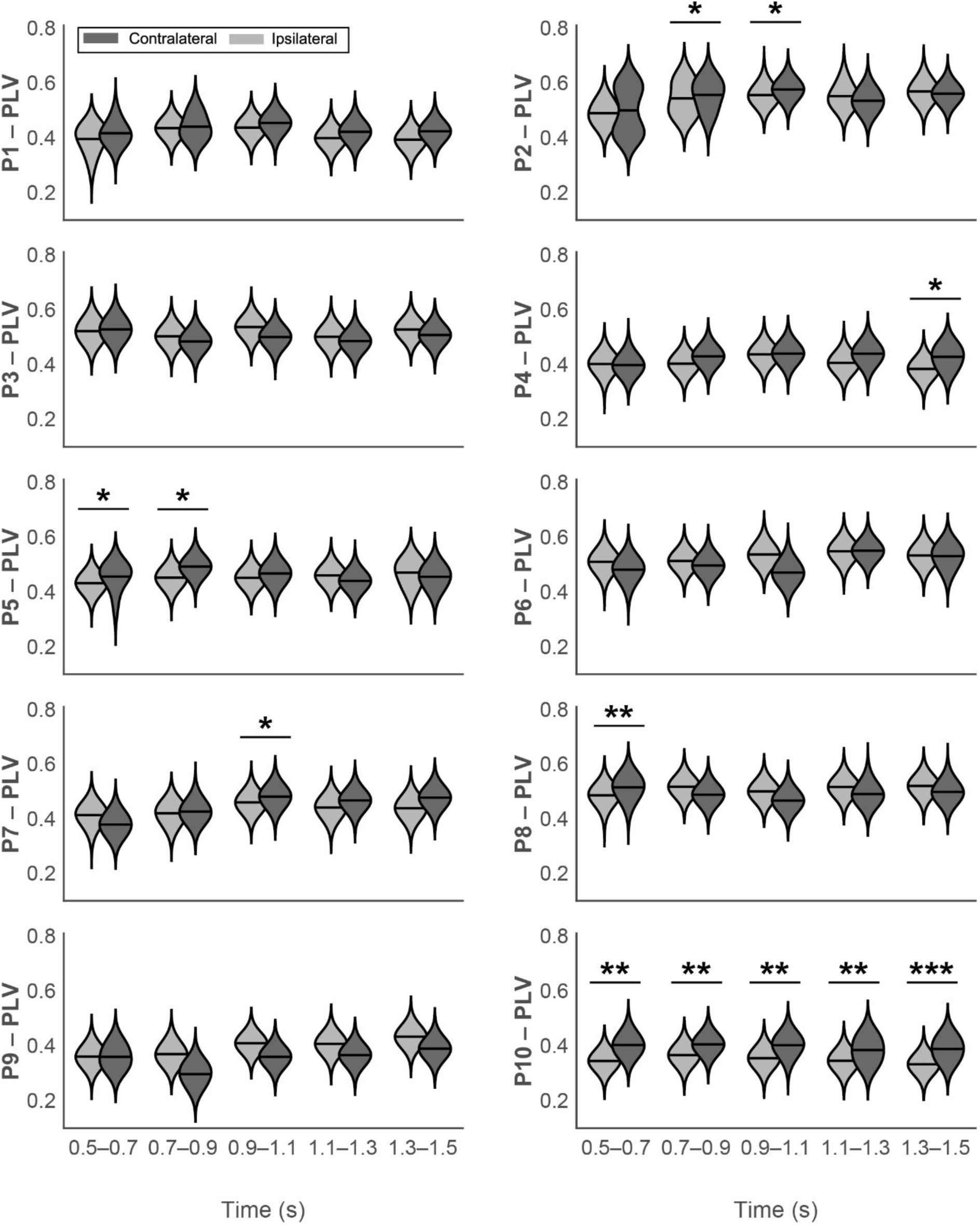
Individual results of upper-alpha cue-locked PLV analysis. Violin plots represent the phase locking values (PLV) averaged over the five time windows (500 to 700, 700 to 900, 1100 to 1300, and 1300 to 1500 ms; t = 0 as cue appearance). Ipsilateral or contralateral scenarios are exhibited as either light grey or dark grey, respectively. \**p* < 0.05, \*\**p* < 0.01.

**Figure 3-2.**
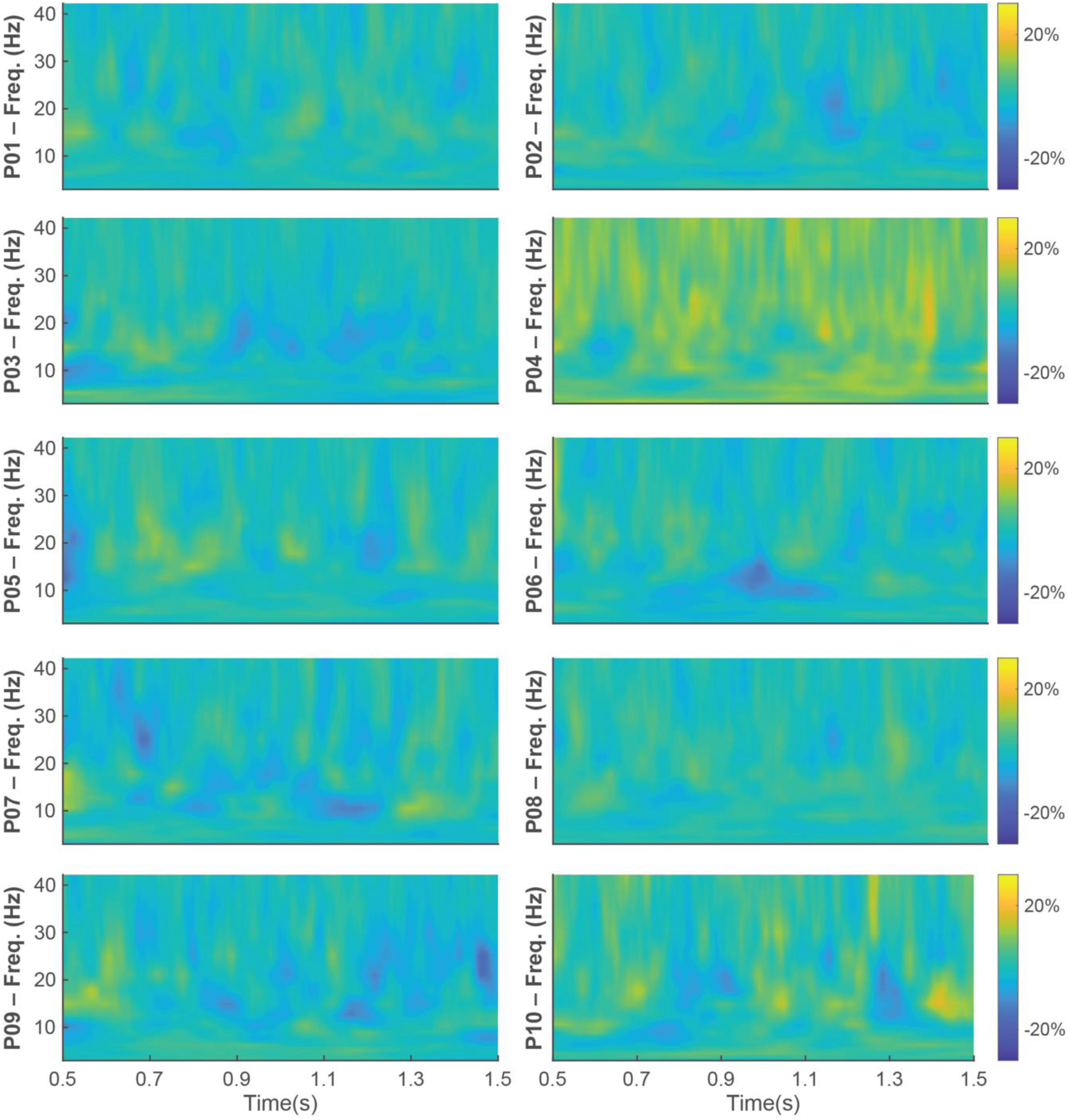
Individual results of cue-locked exploratory PLV analysis. Differences of contra- to ipsilateral PLV are represented over frequencies (2.4 – 42 Hz in 16 logarithmic steps) as a percentage of change regarding the cross-frequency mean of each individual.

**Figure 4-1.**
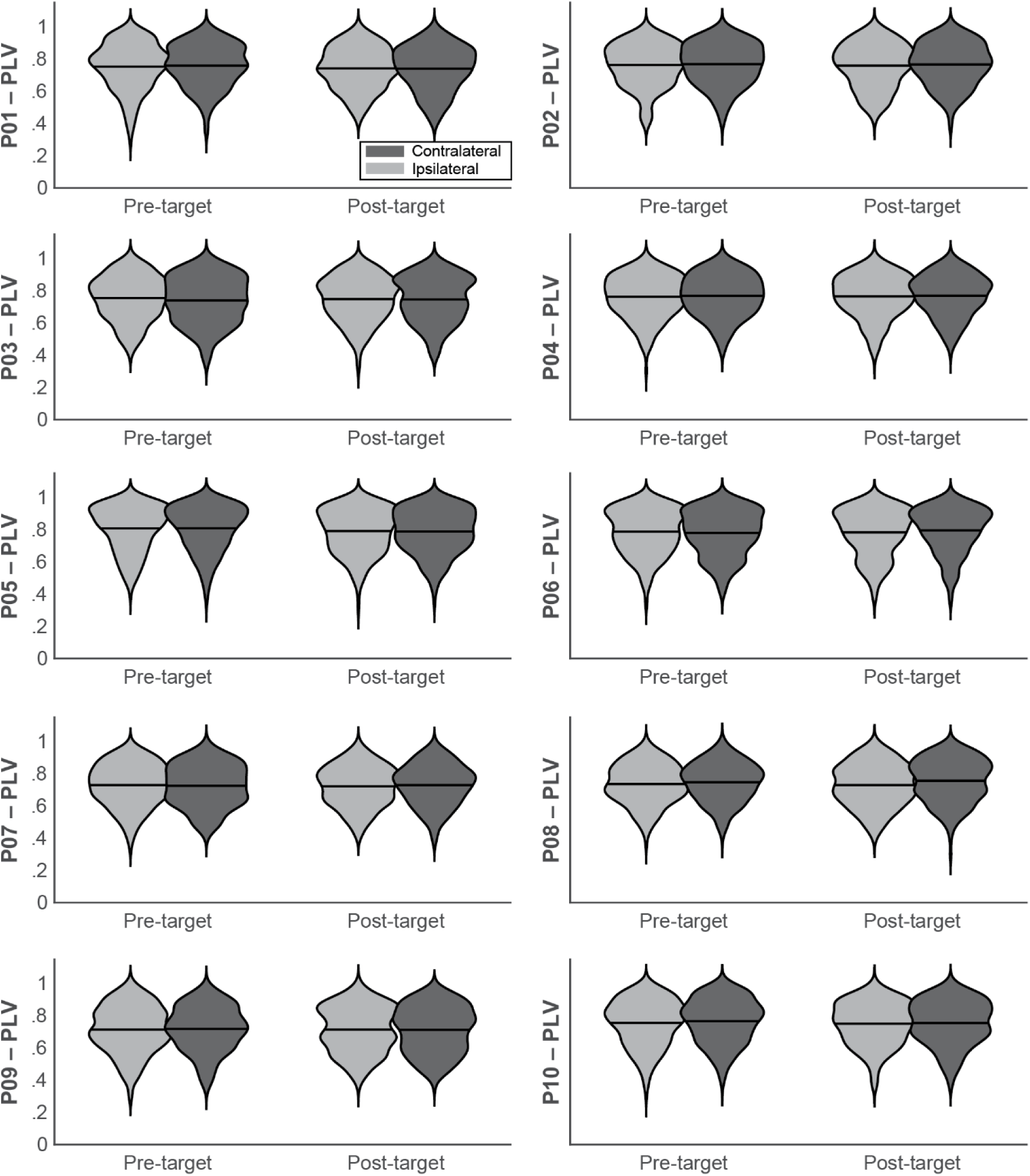
Individual results of target-locked cross-time PLV. Violin plots represent the phase locking values (PLV) obtained by calculating PLV as consistency throughout the pre-target (-200 to 0 ms) and post-target (200 to 400 ms) time windows. Ipsilateral or contralateral scenarios are exhibited as either light grey or dark grey, respectively.

**Figure 4-2.**
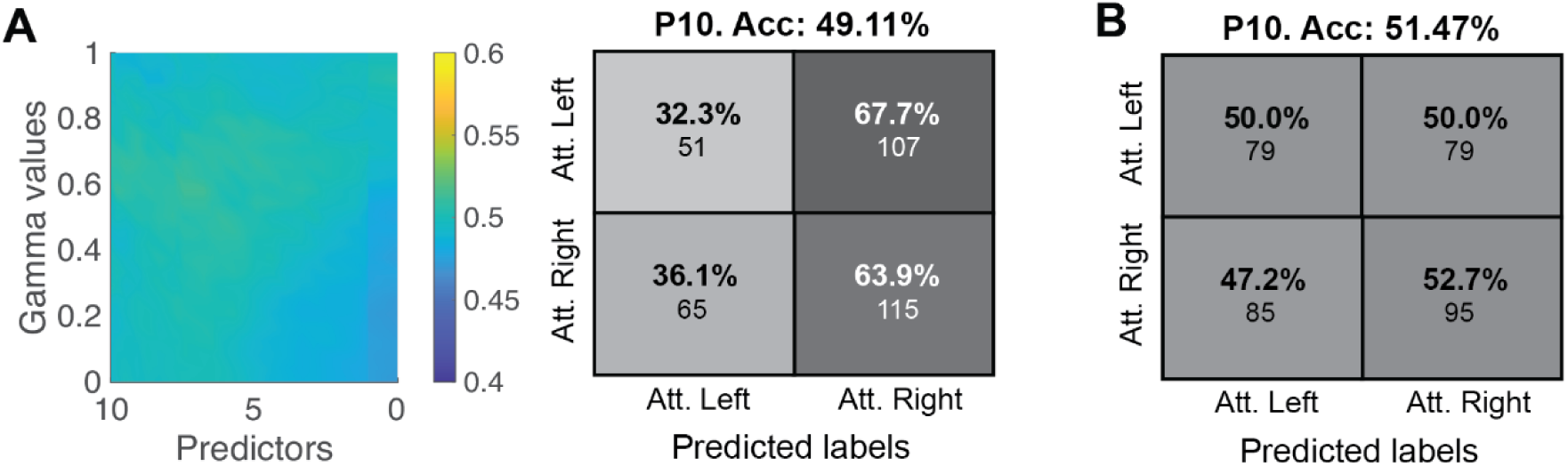
Additional classifier analysis. **(A) Shrinkage linear discriminant analysis**. The leftmost panel reveals how classification error is not modulated by gamma parameter of number of predictors. The rightmost panel presents the confusion matrix. **(B) Riemannian minimum distance to the mean classification results.**

**Figure 5-1.**
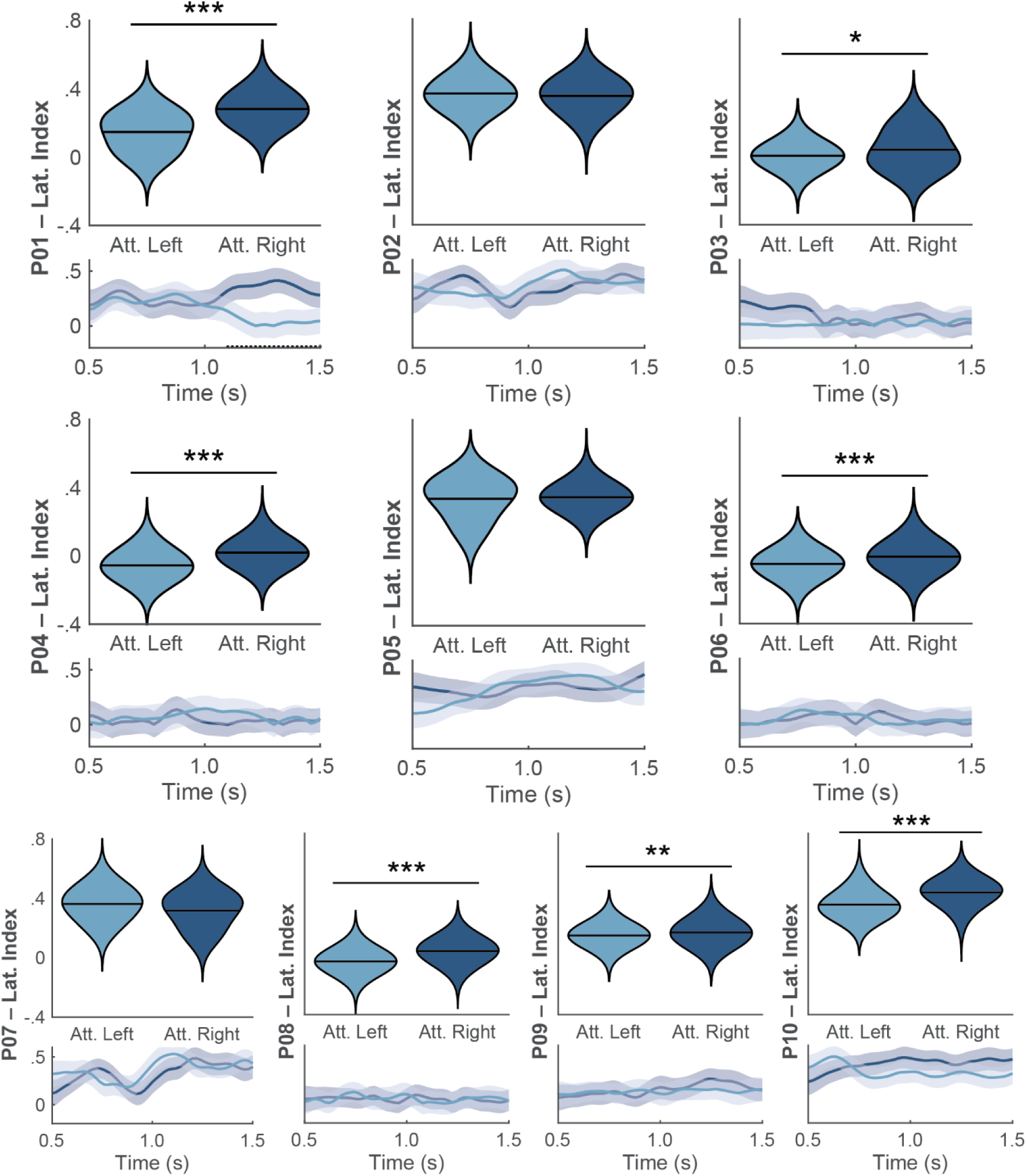
Individual results of lateralisation index. Violin plots represent the averaged lateralised index for *attended left* (light blue) and *attended right* trials (dark blue) over the cue-locked time window. Shaded plots represent lateralisation over time (mean ± SEM). Dots on in the x-axis denote the significant differences over time between *attended left* and *attended right* via cluster-based permutation test. \**p* < 0.05, \*\**p* < 0.01, \*\*\**p* < 0.001.

## REFERENCES

1. Antonov, Plamen A.; Chakravarthi, Ramakrishna; Andersen, Søren K. (2020): Too little, too late, and in the wrong place: Alpha band activity does not reflect an active mechanism of selective attention. In NeuroImage 219, p. 117006. DOI: 10.1016/j.neuroimage.2020.117006.

2. Asplund, Christopher L.; Todd, J. Jay; Snyder, Andy P.; Marois, René (2010): A central role for the lateral prefrontal cortex in goal-directed and stimulus-driven attention. In Nature neuroscience 13 (4), pp. 507–512. DOI: 10.1038/nn.2509.

3. Astrand, Elaine; Enel, Pierre; Ibos, Guilhem; Dominey, Peter Ford; Baraduc, Pierre; Ben Hamed, Suliann (2014a): Comparison of classifiers for decoding sensory and cognitive information from prefrontal neuronal populations. In PloS one 9 (1), e86314. DOI: 10.1371/journal.pone.0086314.

4. Astrand, Elaine; Wardak, Claire; Ben Hamed, Suliann (2014b): Selective visual attention to drive cognitive brain-machine interfaces: from concepts to neurofeedback and rehabilitation applications. In Frontiers in systems neuroscience 8, p. 144. DOI: 10.3389/fnsys.2014.00144.

5. Babiloni, Claudio; Vecchio, Fabrizio; Bultrini, Alessandro; Luca Romani, Gian; Rossini, Paolo Maria (2006): Pre- and poststimulus alpha rhythms are related to conscious visual perception: a high-resolution EEG study. In Cerebral cortex (New York, N.Y. : 1991) 16 (12), pp. 1690–1700. DOI: 10.1093/cercor/bhj104.

6. Baselice, F., Sorriso, A., Rucco, R., & Sorrentino, P. (2018). Phase linearity measurement: A novel index for brain functional connectivity. IEEE transactions on medical imaging, 38(4), 873–882.

7. Benjamini, Yoav; Hochberg, Yosef (1995): Controlling the False Discovery Rate: A Practical and Powerful Approach to Multiple Testing. In Journal of the Royal Statistical Society: Series B (Methodological) 57 (1), pp. 289–300. DOI: 10.1111/j.2517-6161.1995.tb02031.x.

8. Blankertz, Benjamin; Acqualagna, Laura; Dähne, Sven; Haufe, Stefan; Schultze-Kraft, Matthias; Sturm, Irene et al. (2016): The Berlin Brain-Computer Interface: Progress Beyond Communication and Control. In Frontiers in neuroscience 10, p. 530. DOI: 10.3389/fnins.2016.00530.

9. Bonnefond, Mathilde; Kastner, Sabine; Jensen, Ole (2017): Communication between Brain Areas Based on Nested Oscillations. In eNeuro 4 (2). DOI: 10.1523/ENEURO.0153-16.2017.

10. Capotosto, Paolo; Babiloni, Claudio; Romani, Gian Luca; Corbetta, Maurizio (2009): Frontoparietal cortex controls spatial attention through modulation of anticipatory alpha rhythms. In J. Neurosci. 29 (18), pp. 5863–5872. DOI: 10.1523/jneurosci.0539-09.2009.

11. Clayton, Michael S.; Yeung, Nick; Cohen Kadosh, Roi (2018): The many characters of visual alpha oscillations. In The European journal of neuroscience 48 (7), pp. 2498–2508. DOI: 10.1111/ejn.13747.

12. Cohen, Michael X. (2015): Effects of time lag and frequency matching on phase-based connectivity. In Journal of neuroscience methods 250, pp. 137–146. DOI: 10.1016/j.jneumeth.2014.09.005.

13. Cohen, Mike X. (2014): Analysing neural time series data: Theory and practice. Cambridge, Massachusetts, USA: The MIT Press.

14. Corbetta, Maurizio; Shulman, Gordon L. (2002): Control of goal-directed and stimulus-driven attention in the brain. In *Nature reviews*. Neuroscience 3 (3), pp. 201–215. DOI: 10.1038/nrn755.

15. D’Andrea, Antea; Chella, Federico; Marshall, Tom R.; Pizzella, Vittorio; Romani, Gian Luca; Jensen, Ole; Marzetti, Laura (2019): Alpha and alpha-beta phase synchronisation mediate the recruitment of the visuospatial attention network through the Superior Longitudinal Fasciculus. In NeuroImage 188, pp. 722–732. DOI: 10.1016/j.neuroimage.2018.12.056.

16. Doesburg, Sam M.; Bedo, Nicolas; Ward, Lawrence M. (2016): Top-down alpha oscillatory network interactions during visuospatial attention orienting. In NeuroImage 132, pp. 512–519. DOI: 10.1016/j.neuroimage.2016.02.076.

17. Doesburg, Sam M.; Green, Jessica J.; McDonald, John J.; Ward, Lawrence M. (2009): From local inhibition to long-range integration: a functional dissociation of alpha-band synchronisation across cortical scales in visuospatial attention. In Brain research 1303, pp. 97–110. DOI: 10.1016/j.brainres.2009.09.069.

18. Esghaei, Moein; Daliri, Mohammad Reza (2014): Decoding of visual attention from LFP signals of macaque MT. In PloS one 9 (6), e100381. DOI: 10.1371/journal.pone.0100381.

19. Esterman, Michael; Liu, Guanyu; Okabe, Hidefusa; Reagan, Andrew; Thai, Michelle; DeGutis, Joe (2015): Frontal eye field involvement in sustaining visual attention: evidence from transcranial magnetic stimulation. In NeuroImage 111, pp. 542–548. DOI: 10.1016/j.neuroimage.2015.01.044.

20. Foster, Joshua J.; Awh, Edward (2019): The role of alpha oscillations in spatial attention: limited evidence for a suppression account. In Current opinion in psychology 29, pp. 34–40. DOI: 10.1016/j.copsyc.2018.11.001.

21. Foster, Joshua J.; Sutterer, David W.; Serences, John T.; Vogel, Edward K.; Awh, Edward (2017): Alpha-Band Oscillations Enable Spatially and Temporally Resolved Tracking of Covert Spatial Attention. In Psychological science 28 (7), pp. 929–941. DOI: 10.1177/0956797617699167.

22. Foxe, John J.; Snyder, Adam C. (2011): The Role of Alpha-Band Brain Oscillations as a Sensory Suppression Mechanism during Selective Attention. In Frontiers in psychology 2, p. 154. DOI: 10.3389/fpsyg.2011.00154.

23. Fries, Pascal (2005): A mechanism for cognitive dynamics: neuronal communication through neuronal coherence. In Trends in cognitive sciences 9 (10), pp. 474–480. DOI: 10.1016/j.tics.2005.08.011.

24. Fries, Pascal (2015): Rhythms for Cognition: Communication through Coherence. In Neuron 88 (1), pp. 220–235. DOI: 10.1016/j.neuron.2015.09.034.

25. Gaillard, Corentin; Ben Hadj Hassen, Sameh; Di Bello, Fabio; Bihan-Poudec, Yann; VanRullen, Rufin; Ben Hamed, Suliann (2020): Prefrontal attentional saccades explore space rhythmically. In Nature communications 11 (1), p. 925. DOI: 10.1038/s41467-020-14649-7.

26. Green, Jessica J.; McDonald, John J. (2008): Electrical Neuroimaging Reveals Timing of Attentional Control Activity in Human Brain. In PLoS Biol 6 (4), e81. DOI: 10.1371/journal.pbio.0060081.

27. Gregoriou, Georgia G.; Gotts, Stephen J.; Zhou, Huihui; Desimone, Robert (2009): High-frequency, long-range coupling between prefrontal and visual cortex during attention. In Science (New York, N.Y.) 324 (5931), pp. 1207–1210. DOI: 10.1126/science.1171402.

28. Grent-’t-Jong, Tineke; Woldorff, Marty G. (2007): Timing and sequence of brain activity in top-down control of visual-spatial attention. In PLoS Biol 5 (1), e12. DOI: 10.1371/journal.pbio.0050012.

29. Grossmann, A.; Morlet, J. (1984): Decomposition of Hardy Functions into Square Integrable Wavelets of Constant Shape. In SIAM J. Math. Anal. 15 (4), pp. 723–736. DOI: 10.1137/0515056.

30. Haegens, Saskia; Nácher, Verónica; Luna, Rogelio; Romo, Ranulfo; Jensen, Ole (2011): α-Oscillations in the monkey sensorimotor network influence discrimination performance by rhythmical inhibition of neuronal spiking. In Proceedings of the National Academy of Sciences of the United States of America 108 (48), pp. 19377– 19382. DOI: 10.1073/pnas.1117190108.

31. Heilman, K. M.; van den Abell, T. (1980): Right hemisphere dominance for attention: the mechanism underlying hemispheric asymmetries of inattention (neglect). In Neurology 30 (3), pp. 327–330. DOI: 10.1212/wnl.30.3.327.

32. Helfrich, Randolph F.; Fiebelkorn, Ian C.; Szczepanski, Sara M.; Lin, Jack J.; Parvizi, Josef; Knight, Robert T.; Kastner, Sabine (2018): Neural Mechanisms of Sustained Attention Are Rhythmic. In Neuron 99 (4), 854–865.e5. DOI: 10.1016/j.neuron.2018.07.032.

33. Hjorth, B. (1975). An on-line transformation of EEG scalp potentials into orthogonal source derivations. Electroencephalography and clinical neurophysiology, 39(5), 526–530.

34. Hopfinger, J. B.; Buonocore, M. H.; Mangun, G. R. (2000): The neural mechanisms of top-down attentional control. In Nature neuroscience 3 (3), pp. 284–291. DOI: 10.1038/72999.

35. Hung, June; Driver, Jon; Walsh, Vincent (2011): Visual selection and the human frontal eye fields: effects of frontal transcranial magnetic stimulation on partial report analysed by Bundesen’s theory of visual attention. In J. Neurosci. 31 (44), pp. 15904–15913. DOI: 10.1523/jneurosci.2626-11.2011.

36. Jensen, Ole; Bonnefond, Mathilde; Marshall, Tom R.; Tiesinga, Paul (2015): Oscillatory mechanisms of feedforward and feedback visual processing. In Trends in neurosciences 38 (4), pp. 192–194. DOI: 10.1016/j.tins.2015.02.006.

37. Jensen, Ole; Gips, Bart; Bergmann, Til Ole; Bonnefond, Mathilde (2014): Temporal coding organised by coupled alpha and gamma oscillations prioritise visual processing. In Trends in neurosciences 37 (7), pp. 357–369. DOI: 10.1016/j.tins.2014.04.001.

38. Kastner, S.; Ungerleider, L. G. (2000): Mechanisms of visual attention in the human cortex. In Annual review of neuroscience 23, pp. 315–341. DOI: 10.1146/annurev.neuro.23.1.315.

39. Keitel, C., Ruzzoli, M., Dugué, L., Busch, N. A., & Benwell, C. S. (2022). Rhythms in cognition: The evidence revisited. European Journal of Neuroscience, 55(11-12), 2991–3009.

40. Klimesch, Wolfgang (1999): EEG alpha and theta oscillations reflect cognitive and memory performance: a review and analysis. In Brain Research Reviews 29 (2-3), pp. 169–195. DOI: 10.1016/s0165-0173(98)00056-3.

41. Klimesch, Wolfgang (2012): α-band oscillations, attention, and controlled access to stored information. In Trends in cognitive sciences 16 (12), pp. 606–617. DOI: 10.1016/j.tics.2012.10.007.

42. Klimesch, Wolfgang; Sauseng, Paul; Hanslmayr, Simon (2007): EEG alpha oscillations: the inhibition-timing hypothesis. In Brain Research Reviews 53 (1), pp. 63–88. DOI: 10.1016/j.brainresrev.2006.06.003.

43. Lachaux, Jean-Philippe; Rodriguez, Eugenio; Martinerie, Jacques; Varela, Francisco J. (1999): Measuring phase synchrony in brain signals. In Hum. Brain Mapp. 8 (4), pp. 194–208. DOI: 10.1002/(sici)1097-0193(1999)8:4%3C194::aid-hbm4%3E3.0.co;2-c.

44. Lange, Joachim; Oostenveld, Robert; Fries, Pascal (2013): Reduced occipital alpha power indexes enhanced excitability rather than improved visual perception. In J. Neurosci. 33 (7), pp. 3212–3220. DOI: 10.1523/jneurosci.3755-12.2013.

45. Lobier, Muriel; Palva, J. Matias; Palva, Satu (2018): High-alpha band synchronisation across frontal, parietal and visual cortex mediates behavioral and neuronal effects of visuospatial attention. In NeuroImage 165, pp. 222–237. DOI: 10.1016/j.neuroimage.2017.10.044.

46. Lotte, F., Congedo, M., Lécuyer, A., Lamarche, F., & Arnaldi, B. (2007). A review of classification algorithms for EEG-based brain–computer interfaces. Journal of neural engineering, 4(2), R1.

47. Lotte, F., Bougrain, L., Cichocki, A., Clerc, M., Congedo, M., Rakotomamonjy, A., & Yger, F. (2018). A review of classification algorithms for EEG-based brain–computer interfaces: a 10 year update. Journal of neural engineering, 15(3), 031005.

48. Makeig, Scott; Bell, Anthony J.; Jung, Tzyy-Ping; Sejnowski, Terrence J. (1995): Independent Component Analysis of Electroencephalographic Data. In : Proceedings of the 8th International Conference on Neural Information Processing Systems. Cambridge, MA, USA: MIT Press (NIPS’95), pp. 145–151.

49. Maris, Eric; Oostenveld, Robert (2007): Nonparametric statistical testing of EEG- and MEG-data. In Journal of neuroscience methods 164 (1), pp. 177–190. DOI: 10.1016/j.jneumeth.2007.03.024.

50. Marshall, Tom R.; O’Shea, Jacinta; Jensen, Ole; Bergmann, Til O. (2015): Frontal eye fields control attentional modulation of alpha and gamma oscillations in contralateral occipitoparietal cortex. In J. Neurosci. 35 (4), pp. 1638–1647. DOI: 10.1523/jneurosci.3116-14.2015.

51. Meyer, Marlene; Lamers, Didi; Kayhan, Ezgi; Hunnius, Sabine; Oostenveld, Robert (2021): Enhancing reproducibility in developmental EEG research: BIDS, cluster-based permutation tests, and effect sizes. In Developmental cognitive neuroscience 52, p. 101036. DOI: 10.1016/j.dcn.2021.101036.

52. Mostame, Parham; Moharramipour, Ali; Hossein-Zadeh, Gholam-Ali; Babajani-Feremi, Abbas (2019): Statistical Significance Assessment of Phase Synchrony in the Presence of Background Couplings: An ECoG Study. In Brain topography 32 (5), pp. 882–896. DOI: 10.1007/s10548-019-00718-8.

53. Padfield, Natasha; Zabalza, Jaime; Zhao, Huimin; Masero, Valentin; Ren, Jinchang (2019): EEG-Based Brain-Computer Interfaces Using Motor-Imagery: Techniques and Challenges. In Sensors (Basel, Switzerland) 19 (6). DOI: 10.3390/s19061423.

54. Palva, Satu; Palva, J. Matias (2007): New vistas for alpha-frequency band oscillations. In Trends in neurosciences 30 (4), pp. 150–158. DOI: 10.1016/j.tins.2007.02.001.

55. Palva, Satu; Palva, J. Matias (2011): Functional roles of alpha-band phase synchronisation in local and large-scale cortical networks. In Frontiers in psychology 2, p. 204. DOI: 10.3389/fpsyg.2011.00204.

56. Pantazis, Dimitrios; Simpson, Gregory V.; Weber, Darren L.; Dale, Corby L.; Nichols, Thomas E.; Leahy, Richard M. (2009): A novel ANCOVA design for analysis of MEG data with application to a visual attention study. In NeuroImage 44 (1), pp. 164–174. DOI: 10.1016/j.neuroimage.2008.07.012.

57. Pashler, H. (1999): The Psychology of Attention: MIT Press (A Bradford book). Available online at https://books.google.es/books?id=w\_4MyczgUEcC.

58. Petersen, Steven E.; Posner, Michael I. (2012): The attention system of the human brain: 20 years after. In Annual review of neuroscience 35, pp. 73–89. DOI: 10.1146/annurev-neuro-062111-150525.

59. Posner, M. I. (1980): Orienting of attention. In The Quarterly journal of experimental psychology 32 (1), pp. 3–25. DOI: 10.1080/00335558008248231.

60. Rihs, Tonia A.; Michel, Christoph M.; Thut, Gregor (2007): Mechanisms of selective inhibition in visual spatial attention are indexed by alpha-band EEG synchronisation. In The European journal of neuroscience 25 (2), pp. 603–610. DOI: 10.1111/j.1460-9568.2007.05278.x.

61. Ruzzoli, Manuela; Torralba, Mireia; Morís Fernández, Luis; Soto-Faraco, Salvador (2019): The relevance of alpha phase in human perception. In Cortex; a journal devoted to the study of the nervous system and behavior 120, pp. 249–268. DOI: 10.1016/j.cortex.2019.05.012.

62. Sacchet, Matthew D.; LaPlante, Roan A.; Wan, Qian; Pritchett, Dominique L.; Lee, Adrian K. C.; Hämäläinen, Matti et al. (2015): Attention drives synchronisation of alpha and beta rhythms between right inferior frontal and primary sensory neocortex. In J. Neurosci. 35 (5), pp. 2074–2082. DOI: 10.1523/jneurosci.1292-14.2015.

63. Sadaghiani, Sepideh; Kleinschmidt, Andreas (2016): Brain Networks and α-Oscillations: Structural and Functional Foundations of Cognitive Control. In Trends in cognitive sciences 20 (11), pp. 805–817. DOI: 10.1016/j.tics.2016.09.004.

64. Samaha, Jason; Bauer, Phoebe; Cimaroli, Sawyer; Postle, Bradley R. (2015): Top-down control of the phase of alpha-band oscillations as a mechanism for temporal prediction. In Proceedings of the National Academy of Sciences of the United States of America 112 (27), pp. 8439–8444. DOI: 10.1073/pnas.1503686112.

65. Sauseng, P.; Klimesch, W.; Stadler, W.; Schabus, M.; Doppelmayr, M.; Hanslmayr, S. et al. (2005): A shift of visual spatial attention is selectively associated with human EEG alpha activity. In The European journal of neuroscience 22 (11), pp. 2917– 2926. DOI: 10.1111/j.1460-9568.2005.04482.x.

66. Siegel, Markus; Donner, Tobias H.; Oostenveld, Robert; Fries, Pascal; Engel, Andreas K. (2008): Neuronal synchronisation along the dorsal visual pathway reflects the focus of spatial attention. In Neuron 60 (4), pp. 709–719. DOI: 10.1016/j.neuron.2008.09.010.

67. Silvanto, Juha; Lavie, Nilli; Walsh, Vincent (2006): Stimulation of the human frontal eye fields modulates sensitivity of extrastriate visual cortex. In Journal of neurophysiology 96 (2), pp. 941–945. DOI: 10.1152/jn.00015.2006.

68. Simpson, Gregory V.; Weber, Darren L.; Dale, Corby L.; Pantazis, Dimitrios; Bressler, Steven L.; Leahy, Richard M.; Luks, Tracy L. (2011): Dynamic activation of frontal, parietal, and sensory regions underlying anticipatory visual spatial attention. In J. Neurosci. 31 (39), pp. 13880–13889. DOI: 10.1523/jneurosci.1519-10.2011.

69. Thatcher, R. W., Soler, E. P., North, D. M., & Otte, G. (2020). Independent Components Analysis “Artifact Correction” Distorts EEG Phase in Artifact Free Segments. J Neurol Neurobiol, 6(4), 5–7.

70. Thut, Gregor; Nietzel, Annika; Brandt, Stephan A.; Pascual-Leone, Alvaro (2006): Alpha-band electroencephalographic activity over occipital cortex indexes visuospatial attention bias and predicts visual target detection. In The Journal of neuroscience : the official journal of the Society for Neuroscience 26 (37), pp. 9494– 9502. DOI: 10.1523/JNEUROSCI.0875-06.2006.

71. Tonin, L.; Leeb, R.; Del R Millán, J. (2012): Time-dependent approach for single trial classification of covert visuospatial attention. In Journal of neural engineering 9 (4), p. 45011. DOI: 10.1088/1741-2560/9/4/045011.

72. Tonin, L.; Leeb, R.; Sobolewski, A.; Del Millán, J. R. (2013): An online EEG BCI based on covert visuospatial attention in absence of exogenous stimulation. In Journal of neural engineering 10 (5), p. 56007. DOI: 10.1088/1741-2560/10/5/056007.

73. Treder, Matthias S.; Bahramisharif, Ali; Schmidt, Nico M.; van Gerven, Marcel A. J.; Blankertz, Benjamin (2011): Brain-computer interfacing using modulations of alpha activity induced by covert shifts of attention. In Journal of neuroengineering and rehabilitation 8, p. 24. DOI: 10.1186/1743-0003-8-24.

74. Tremblay, Sébastien; Doucet, Guillaume; Pieper, Florian; Sachs, Adam; Martinez-Trujillo, Julio (2015): Single-Trial Decoding of Visual Attention from Local Field Potentials in the Primate Lateral Prefrontal Cortex Is Frequency-Dependent. In J. Neurosci. 35 (24), pp. 9038–9049. DOI: 10.1523/jneurosci.1041-15.2015.

75. van Diepen, Rosanne M.; Foxe, John J.; Mazaheri, Ali (2019): The functional role of alpha-band activity in attentional processing: the current zeitgeist and future outlook. In Current opinion in psychology 29, pp. 229–238. DOI: 10.1016/j.copsyc.2019.03.015.

76. van Gerven, Marcel; Jensen, Ole (2009): Attention modulations of posterior alpha as a control signal for two-dimensional brain-computer interfaces. In Journal of neuroscience methods 179 (1), pp. 78–84. DOI: 10.1016/j.jneumeth.2009.01.016.

77. VanRullen, Rufin (2016): Perceptual Cycles. In Trends in cognitive sciences 20 (10), pp. 723–735. DOI: 10.1016/j.tics.2016.07.006.

78. Veniero, D., Gross, J., Morand, S., Duecker, F., Sack, A. T., & Thut, G. (2021). Top-down control of visual cortex by the frontal eye fields through oscillatory realignment. Nature communications, 12(1), 1–13.

79. Vigué-Guix, I., Moris Fernandez, L., Torralba Cuello, M., Ruzzoli, M., & Soto-Faraco, S. (2022). Can the occipital alpha-phase speed up visual detection through a real-time EEG-based brain–computer interface (BCI)?. European Journal of Neuroscience, 55(11-12), 3224–3240.

80. Yamagishi, Noriko; Callan, Daniel E.; Goda, Naokazu; Anderson, Stephen J.; Yoshida, Yoshikazu; Kawato, Mitsuo (2003): Attentional modulation of oscillatory activity in human visual cortex. In NeuroImage 20 (1), pp. 98–113. DOI: 10.1016/s1053-8119(03)00341-0.

81. Zago, Laure; Petit, Laurent; Jobard, Gael; Hay, Julien; Mazoyer, Bernard; Tzourio-Mazoyer, Nathalie et al. (2017): Pseudoneglect in line bisection judgement is associated with a modulation of right hemispheric spatial attention dominance in right-handers. In Neuropsychologia 94, pp. 75–83. DOI: 10.1016/j.neuropsychologia.2016.11.024.

